# A CK2α–G3BP1 signaling axis regulates local translation in developing neurons and is disrupted in OCNDS

**DOI:** 10.64898/2026.08.11.744218

**Authors:** Manasi Agrawal, Meghal Desai, Shruti Ghumra, Yashashree Bhorkar, Brandon J. Vaglio, Kyle Stokes, Krishna Rana, Ny-Ziah Hamilton Hill, Peace Nweze, Nethra Sriram, Alana LoRé, Riki Kawaguchi, Bonnie L. Firestein, Jack Parent, Daniel H. Geschwind, Heike Rebholz, Pabitra K. Sahoo

**Affiliations:** Department of Biological Sciences, Rutgers University – Newark, Newark, NJ, USA; Department of Cell Biology and Neuroscience, Rutgers, The State University of New Jersey, Piscataway, NJ, USA; Biomedical Engineering Program, Rutgers, The State University of New Jersey, Piscataway, NJ, USA; Department of Neurology, University of Michigan, Ann Arbor, MI, USA; Department of Chemistry and Environmental Sciences, New Jersey Institute of Technology, Newark, NJ, USA; Center for Neurobehavioral Genetics, Semel Institute for Neuroscience and Human Behavior, University of California, Los Angeles, Los Angeles, CA 90095, USA; Michigan Neuroscience Institute, Ann Arbor, MI, USA; VA Ann Arbor Health System, Ann Arbor, MI, USA; Program in Neurogenetics, Departments of Neurology and Human Genetics, Institute of Precision Health, David Geffen School of Medicine, University of California, Los Angeles, Los Angeles, CA, USA; Center for Autism Research and Treatment, Department of Psychiatry and Semel Institute, University of California, Los Angeles, Los Angeles, CA, USA; Department of Neurology, University of California, Los Angeles, Los Angeles, CA, USA; Université Paris Cité, Institute of Psychiatry and Neuroscience of Paris (IPNP), INSERM U1266, Laboratory of Signaling mechanisms in neurological disorders, 75014 Paris, France; GHU-Paris Psychiatrie et Neurosciences, Hôpital Sainte-Anne, F-75014 Paris, France

**Keywords:** Local Protein Synthesis, G3BP1, CK2α, Stress Granules, Neurodevelopment, OCNDS, Liquid-Liquid Phase Separation, RNA Granules

## Abstract

Neurodevelopmental disorders are frequently caused by mutations in pleiotropic kinases, yet downstream effectors driving neuronal pathology remain undefined. Here, we identify the G3BP1-dependent stress granule pathway as the dominant effector of casein kinase 2 (CK2α) in developing neurons, implying that its dysregulation underlies the neurodevelopmental deficits of Okur-Chung neurodevelopmental syndrome (OCNDS). OCNDS-associated CK2α mutations reduce phosphorylation of G3BP1 at serine 149, promoting aberrant phase separation and persistent granules that sequester neuronal mRNAs and suppress local protein synthesis across axonal and dendritic compartments. These phenotypes produce allele-specific deficits in neuronal morphogenesis, synaptic abundance, and network excitability, which are conserved in a knock-in mouse model and in patient-derived iPSC neurons. *G3bp1* knockdown rescues translational and morphological phenotypes across all OCNDS alleles, demonstrating that restoring granule homeostasis reverses neuronal pathology. Together, these findings establish OCNDS as a disorder of compartment-specific translational dysregulation driven by impaired CK2α–G3BP1 control of RNA granule homeostasis.

**Summary:** OCNDS mutations disrupt CK2α–G3BP1 signaling, causing persistent granules and defective neuronal translation and development.

## INTRODUCTION

Okur-Chung neurodevelopmental syndrome (OCNDS) is a rare autosomal dominant disorder caused by *de novo* mutations in *CSNK2A1*, the gene encoding the catalytic alpha subunit of casein kinase 2 (CK2α)(*1–3*). OCNDS is characterized by a broad clinical spectrum that includes intellectual disability, autism spectrum disorder, language impairment, hypotonia, and global developmental delay(*4, 5*). Since the identification of OCNDS-associated *CSNK2A1* mutations through exome sequencing, more than one hundred *CSNK2A1* variants have been reported, highlighting the genetic and phenotypic heterogeneity of the disorder(*6, 7*). Despite this growing mutational catalog, the cellular and molecular mechanisms by which individual CK2α mutations disrupt neuronal development remain poorly understood, posing a critical barrier to the development of targeted therapies.

CK2 is a highly conserved, ubiquitously expressed serine/threonine kinase that functions as a tetrameric holoenzyme composed of two catalytic subunits (CK2α and CK2α’) and two regulatory CK2β subunits(*8, 9*). It is constitutively active and regulates diverse cellular processes, including embryonic development, cell cycle progression, and cellular stress adaptation(*9, 10*). Consistent with its role in embryonic development, genetic disruption of CK2 subunits is embryonically lethal in mice(*11, 12*). CK2 is highly expressed in the nervous system and has been implicated in receptor trafficking and synaptic signaling(*13–16*) as well as in neurodegenerative diseases, such as Alzheimer’s and Parkinson’s(*17–19*). With nearly five hundred known substrates, CK2 occupies a central node in cellular signaling networks, and perturbations in its activity have the potential to simultaneously disrupt multiple downstream pathways(*20, 21*).

Biochemical characterization of OCNDS-associated CK2α variants has demonstrated that disease mutations variably reduce kinase activity toward canonical substrates, with certain alleles, such as K198R, the most prevalent variant in OCNDS patients, additionally altering substrate specificity(*5, 22–24*). Notably, CK2α protein levels are preserved in a subset of patient-derived cells studied thus far and the *CSNK2A1^K198R/+^* knock-in mouse model, indicating that disease phenotypes arise from alterations in kinase function rather than haploinsufficiency(*4, 23–25*). OCNDS follows an autosomal dominant inheritance pattern(*2*). Many disease-associated mutants of CK2α retain the ability to assemble into holoenzyme complexes(*26*). As a result, these mutant proteins may exert dominant-negative effects by competing with the wild-type kinase for access to substrates or for binding to regulatory subunits. Taken together, these features predict that different *CSNK2A1* mutations will disrupt overlapping but non-identical sets of downstream pathways. Identifying the most functionally dominant effectors of CK2α activity in neurons is essential for understanding disease pathogenesis.

RasGAP SH3 domain binding protein 1 (G3BP1) is an RNA-binding protein that serves as a core nucleator of stress granules (SGs), dynamic condensates that generally form in response to cellular stress, and sequester mRNAs for protection and temporary storage(*27–31*). In neurons, G3BP1 granules function as compartment-specific translational checkpoints that modulate the availability of mRNAs required for neuronal growth, morphogenesis, and homeostasis(*21, 27, 29, 32, 33*). Due to the extreme polarity of neurons, neuronal development and function require precise spatial and temporal regulation of mRNA translation in distal axons and dendrites(*34–39*). Recent work from our laboratory has demonstrated that disassembly of G3BP1 granules is a key mechanism for releasing axonally localized mRNAs for local translation(*27–29*). Promoting this disassembly enhances axon regeneration in both the peripheral and central nervous system neurons. Critically, G3BP1 is a direct substrate of CK2α, which phosphorylates G3BP1 at serine 149 (G3BP1^PS149^) to raise the RNA concentration threshold required for granule nucleation and thereby promote granule disassembly(*20, 28, 40*). After sciatic nerve axotomy, local upregulation of CK2α via axonal translation in injured axons drives G3BP1^PS149^-dependent granule disassembly and mRNA release, linking kinase activity to translational derepression in a spatially regulated manner(*28*). Whether this CK2α–G3BP1 axis operates during neuronal development and whether its dysregulation by OCNDS-associated CK2α mutations contributes to neurodevelopmental deficits remains uninvestigated.

Here, we demonstrate that OCNDS-associated CK2α mutations reduce G3BP1^PS149^ phosphorylation and drive compartment-specific accumulation of persistent, less dynamic SGs in developing cortical neurons. These granules sequester neurodevelopmentally critical mRNAs and broadly suppress local protein synthesis across axonal and dendritic compartments. We show that these phenotypes are conserved in a *CSNK2A1^K198R/+^*knock-in mouse model and in iPSC-derived neurons from an OCNDS patient with the *CSNK2A1^K198R^* mutation. We find CK2α mutation-specific deficits in neuronal morphogenesis, synapse formation, and network activity. Crucially, siRNA-mediated knockdown of *G3bp1* rescues both local translation and the full spectrum of morphological and synaptic phenotypes across all OCNDS-associated CK2α mutants tested. Together, these data establish dysregulated G3BP1-dependent translational control as the central mechanism linking CK2α loss-of-function to neurodevelopmental pathology in OCNDS and demonstrate that restoring granule homeostasis rescues the neuronal and synaptic deficits caused by disease-associated CK2*α* mutants.

## RESULTS

### Expression of OCNDS-associated CK2α mutants results in larger and more persistent stress granules (SGs)

SGs are dynamic ribonucleoprotein assemblies that regulate mRNA storage and repress mRNA translation, with key scaffold proteins G3BP1 and G3BP2 recruiting additional RNA-binding proteins, such as CAPRIN1, to drive SG assembly(*41, 42*). Phosphorylation of G3BP1^PS149^ by CK2α increases G3BP1 mRNA binding threshold for phase separation(*43*). Previously, we demonstrated that in the peripheral nervous system, injury to the sciatic nerve increases the local translation of *Csnk2a1* mRNA, and the nascent CK2α protein, in turn, phosphorylates G3BP1^S149^ to release the axonal mRNAs for translation(*28*). Since OCNDS-associated *CSNK2A1* mutations are autosomal dominant and variably reduce CK2α kinase activity(*24*), we asked if expression of these mutants would affect SG dynamics.

We find that expression of OCNDS-associated CK2α^R47G^, CK2α^R47Q^, CK2α^K198R^, and CK2α^R312W^ mutants, but not of CK2α^WT^, in COS-7 cells leads to SG formation even in the absence of any external stressors (**Fig. 1A-B**, **Fig. S1**). In the presence of sodium arsenite stress, the COS-7 cells expressing the CK2α^R47G^ and CK2α^K198R^ mutants show a significant increase in the number of SGs per cell (**Fig. 1A, Fig. S1**). Even after stress removal, a higher percentage of COS-7 cells expressing the CK2α mutants show significantly increased numbers of SGs per cell, suggesting that the OCNDS-associated CK2α mutants can inhibit SG disassembly (**Fig. 1A-B, Fig. S1**). These findings are consistent with a previous study that inhibited CK2α in melanoma cells, leading to enhanced SG formation during recovery after sodium arsenite stress(*40*).

**Figure 1:**
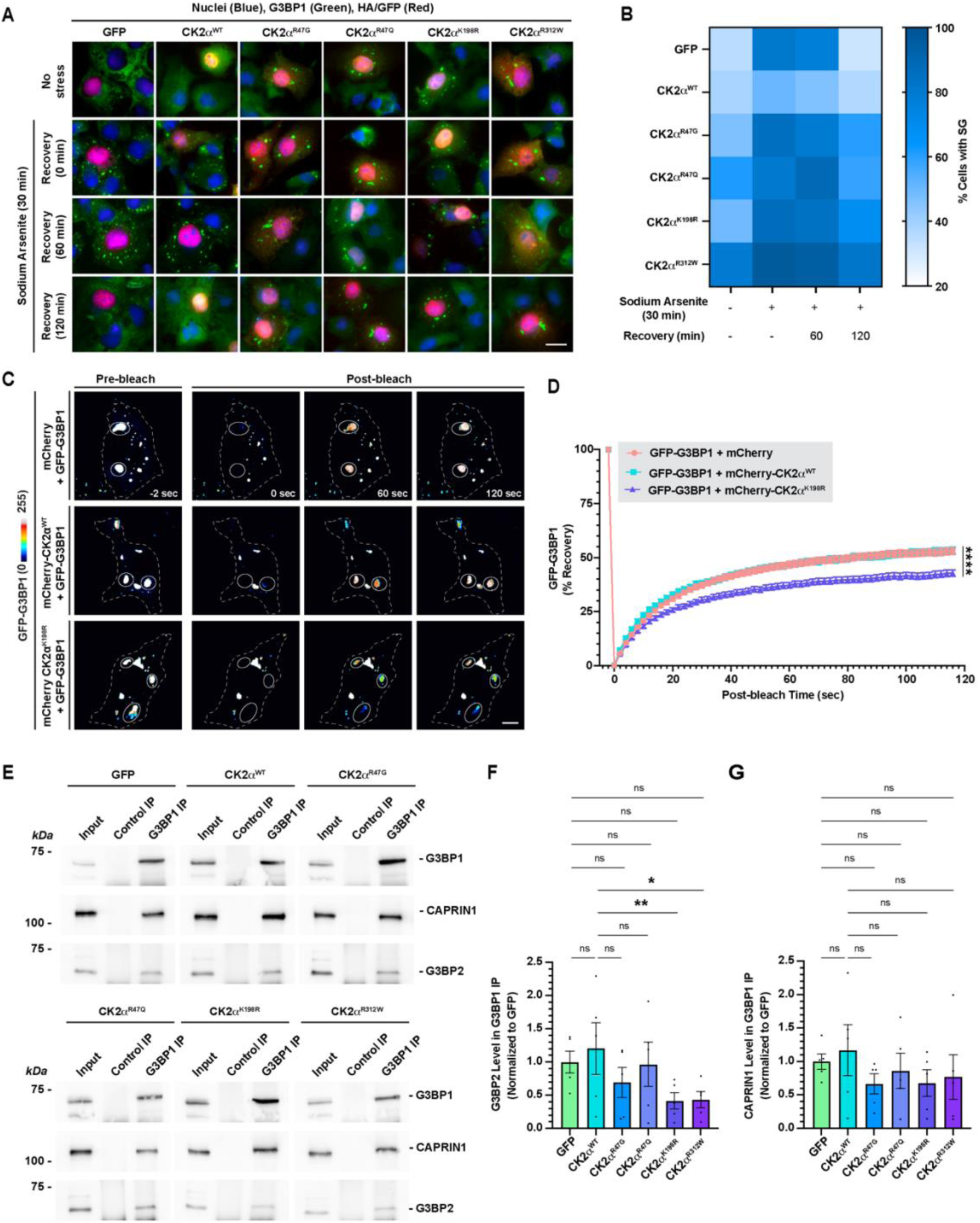
OCNDS-associated CK2α mutants inhibit stress granule disassembly. ***A,*** Representative epifluorescent images of COS-7 cells transfected with either GFP, HA tagged CK2α^WT^ or CK2α mutants, immunostained for G3BP1, HA, and DAPI are shown. Cells were treated with 0.5 mM sodium arsenite for 30 min followed by recovery (60 min and 120 min) from stress. (Scale bar, 20 µm). ***B,*** Quantification of percentage cells with SGs is shown in the heatmap as the mean across three biological replicates. ***C,*** Representative FRAP image sequences for COS-7 cells co-transfected with either mCherry, mCherry-CK2α^WT^, or mCherry-CK2α^K198R^ and GFP-G3BP1 constructs are shown. Dotted lines represent the cell boundary, and circles represent the photobleached ROIs for SGs. Videos for these images are included in the Supplementary Movie 1 (Scale bar, 10 μm). ***D,*** Quantification of FRAP assays in **(C)** is shown as the percent recovery of GFP-G3BP1 signal, with mean ± SEM. n ≥ 150 SGs (3-5 SGs per cell) across three biological replicates; \*\*\*\**p* ≤ 0.0001 between mCherry or mCherry-CK2α^WT^ and mCherry-CK2α^K198R^ by two-way ANOVA with Tukey’s multiple comparisons. ***E,*** Immunoblots of G3BP1 IP from HEK293T cells transfected with GFP control or HA-tagged CK2α constructs are shown. The blots were probed with antibodies against G3BP1, CAPRIN1, and G3BP2. ***F-G*,** Densitometric quantifications of G3BP2 and CAPRIN1 co-immunoprecipitated with G3BP1 and normalized to the GFP control group are shown as mean ± SEM across five biological replicates; *p ≤ 0.05, **p ≤ 0.01 by Friedman test with Dunn’s multiple comparisons.

As the K198R mutation is the most prevalent in OCNDS patients(*22*), we tested whether expression of this mutant affects SG dynamicity. Fluorescence recovery after photobleaching (FRAP) assay shows that COS-7 cells expressing the CK2α^K198R^ mutant display significantly reduced G3BP1 granule dynamics (**Fig. 1C-D, Movie S1**). Both G3BP1 and its paralog, G3BP2, are necessary and sufficient to form the core of SGs(*44–46*). CAPRIN1 regulates liquid-liquid phase separation (LLPS) by interaction with G3BP1(*41, 42*). To assess whether CK2α mutations also alter the molecular composition of the SG core, we used HEK293T cells, which offer higher transfection efficiency and protein yield than COS-7 cells, making them better suited for co-immunoprecipitation studies. Our results show that expression of all OCNDS-associated CK2α mutants results in a trend toward reduced interaction between G3BP1 and G3BP2, and between G3BP1 and CAPRIN1. The CK2α^K198R^ and CK2α^R312W^ mutants show a significant reduction in the interaction of G3BP1 with G3BP2 (**Fig. 1E-G**). Overall, our data indicate that expression of the kinase-defective OCNDS-associated CK2α mutants results in persistent, less dynamic SGs with defective SG cores.

### OCNDS-associated CK2α mutants differentially affect neuronal development and maturation in a compartment-specific manner

Most of the OCNDS-associated CK2α mutants have not yet been tested to determine exactly how they impair neurodevelopment at the cellular level. To test this, we expressed the CK2α^WT^, CK2α^R47G^, CK2α^R47Q^, CK2α^K198R^, and CK2α^R312W^ mutants or GFP in cultured rat cortical neurons and assessed neuronal growth, maturation, and activity over different time points (**Fig. 2A**). To analyze the axonal and dendritic growth defects, we used 11 *DIV* neurons. Neurons expressing CK2α^R47G^, CK2α^K198R^, and CK2α^R312W^ mutants exhibit reduced axonal growth, while CK2α^R47Q^, CK2α^K198R^, and CK2α^R312W^ show increased axonal branching (**Fig. 2B-D**). Interestingly, while CK2α^R47Q^ increases dendritic growth, the CK2α^K198R^ and CK2α^R312W^ mutants reduce dendritic growth (**Fig. 2B, E-F**). None of the mutants affected dendritic branching (**Fig. 2B, E-F**). Overall, these findings suggest that OCNDS-associated CK2α mutants differentially impair neuronal morphology in a compartment-specific manner.

**Figure 2:**
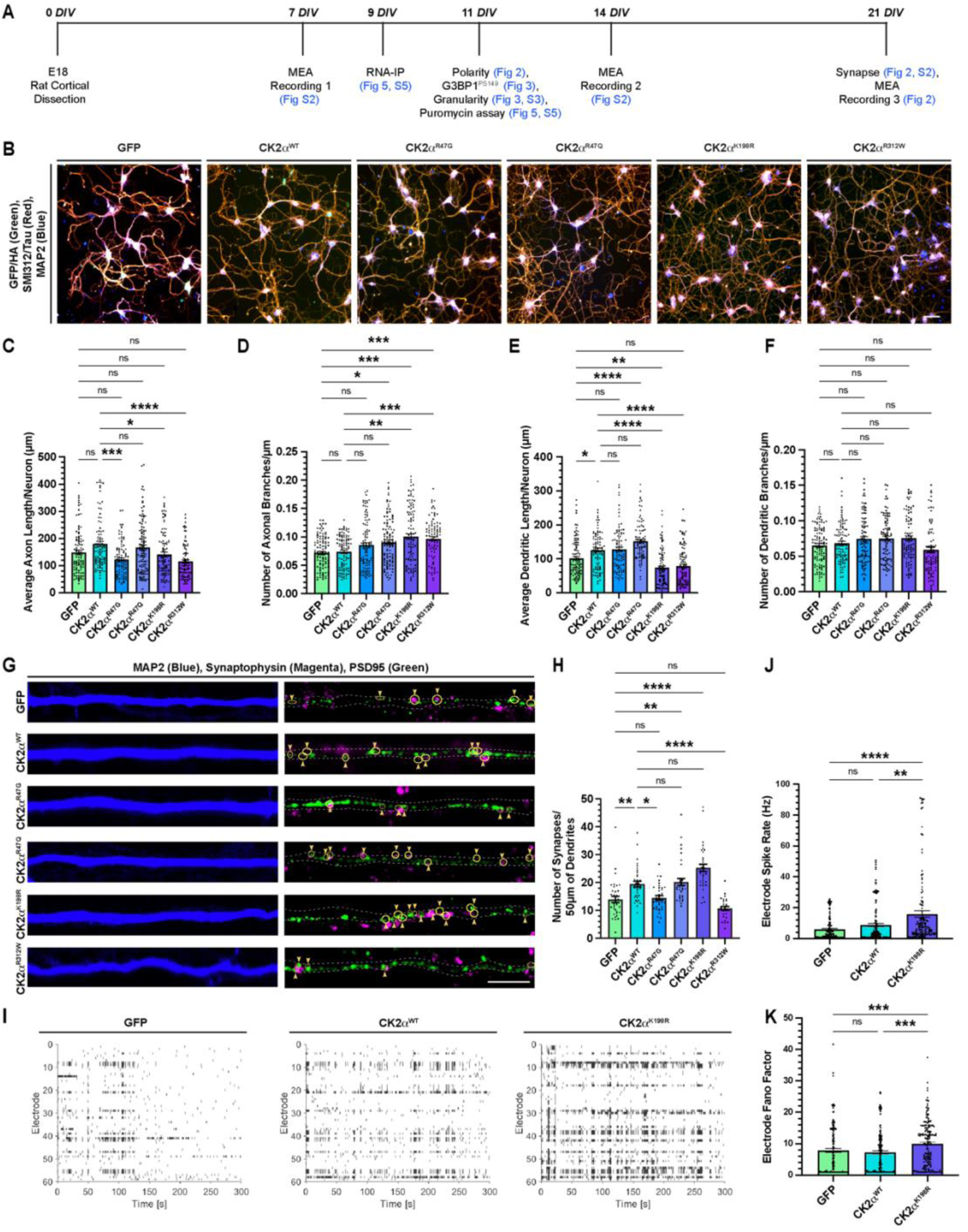
OCNDS-associated CK2α mutants alter developing cortical neuronal growth and synaptogenesis. ***A,*** Timeline for primary rat cortical neurons cultured for various analyses is shown. ***B,*** Representative epifluorescence images of GFP/HA, SMI312/Tau, and MAP2 immunostained E18 rat cortical neurons transfected with the indicated constructs are shown (Scale bar, 100 µm). ***C-F,*** Quantifications of average axonal length **(C)**, branching complexity of axons **(D)**, average dendritic length **(E)**, and branching complexity of dendrites **(F)** using WIS Neuromath are shown as mean ± SEM. n ≥ 88 neurons across three biological replicates; *p ≤ 0.05, **p ≤ 0.01, ***p ≤ 0.001, ****p ≤ 0.0001 by Kruskal-Wallis test with Dunn’s multiple comparison. ***G,*** Representative confocal images of embryonic rat cortical neurons expressing either GFP, CK2α^WT^, or CK2α mutants cultured for 21 *DIV* and immunostained for MAP2, Synaptophysin, PSD95, and GFP or HA are shown (Scale bar, 5 µm). ***H,*** Quantification of the number of synapses is shown as mean ± SEM. n ≥ 30 neurites; *p ≤ 0.05, **p ≤ 0.01, ****p ≤ 0.0001 by Kruskal-Wallis test with Dunn’s multiple comparisons. ***I-K,*** Raster plots for activity detected using MEAs for embryonic rat cortical neurons expressing either GFP, CK2α^WT^, or CK2α^K198R^ cultured for 21 *DIV* are shown (**I**). Quantifications of electrode spike rate **(J)** and electrode Fano factor **(K)** are shown as mean ± SEM. n = recording from 177 electrode across three technical replicates; **p ≤ 0.01, ***p ≤ 0.001, ****p ≤ 0.0001 by Kruskal-Wallis test with Dunn’s multiple comparison.

To assess the effect of mutations on synapse formation, we focused on neurons cultured for 21 *DIV*. Neurons overexpressing CK2α^WT^ exhibit a significantly higher number of synapses than control GFP-expressing neurons (**Fig. 2G, H**), as previously reported(*23*). In contrast, neurons expressing CK2α^K198R^ display a significant increase in the number of synapses compared to the CK2α^WT^-expressing neurons (**Fig. 2G, H**). Conversely, neurons expressing CK2α^R47G^ and CK2α^R312W^ show a reduced number of synapses (**Fig. 2G, H**). There is no significant change in the number of individual PSD95 and synaptophysin puncta compared to the GFP- or CK2α^WT^-expressing neurons (**Fig. S2A-B**). Consistent with our data, the OCNDS mouse model carrying the *CSNK2A1^K198R^*mutation shows increased synaptic numbers in hippocampal neurons(*23*).

Next, we examined how increased synapse number in the CK2α^K198R^ mutant affects spontaneous neuronal activity using microelectrode array (MEA) analysis at 7, 14, and 21 *DIV*. At 7 *DIV*, while both electrode spiking rate and Fano Factor (which detects the variability of neural spiking over time) are significantly lower in CK2α^WT^-expressing neurons than in the GFP-expressing control neurons, these measures are significantly higher in the CK2α^K198R^ mutant-expressing neurons (**Fig. S2C, E-F**). By 14 *DIV*, the electrode spike rate in both CK2α^WT^- and CK2α^K198R^-mutant-expressing neurons is significantly higher than in the GFP-expressing control (H = 22.64; p ≤ 0.0001) (**Fig. S2F, G)**. The Electrode Fano Factor for the CK2α^K198R^ mutant is not statistically different from controls at 14 *DIV* (H = 3.170; p = 0.2025) (**Fig. S2G**). At 21 *DIV*, CK2α^WT^-expressing neurons do not differ significantly from the GFP-expressing control in spike rate, whereas neurons expressing the CK2α^K198R^ mutant show significantly higher spike rates (H = 22.45; p ≤ 0.0001) and higher Fano Factors (H = 20.82; p ≤ 0.0001) than both CK2α^WT^- and the GFP-expressing control, indicating that detected spikes not only increase but also exhibit a more irregular firing pattern (**Fig. 2I-K**). Together, these findings suggest that CK2α kinase activity bidirectionally regulates the maturation of spontaneous network activity, with altered CK2α function driving both excitatory synaptic overabundance and increased and irregular neuronal firing.

### OCNDS-associated CK2α mutants reduce G3BP1 phosphorylation at S149 and increase SG assembly

CK2α is part of the CK2 holoenzyme, which is a serine threonine kinase with nearly 500 substrates, including G3BP1(*20, 28, 47*). CK2α phosphorylates G3BP1 at S149(*20, 28, 40*), and we have previously shown that this leads to G3BP1 granule disassembly and the release of axonal mRNAs in peripheral sensory neurons(*28*). To determine whether CK2α mutations associated with OCNDS alter the phosphorylation status of G3BP1^PS149^, we examined G3BP1^PS149^ levels in cultured developing rat cortical neurons expressing GFP, CK2α^WT^, CK2α^R47G^, CK2α^R47Q^, CK2α^K198R^, or CK2α^R312W^. G3BP1^PS149^ levels are higher in CK2α^WT^-expressing neurons compared to GFP-expressing controls across all neuronal compartments, including the soma, axons, and dendrites (**Fig. 3A-D**). Among the OCNDS-associated mutants, CK2α^R47Q^ similarly elevates G3BP1^PS149^ in the soma, while all other mutants fail to do so (**Fig. 3A-B**). In both axons and dendrites, all four mutants show a significant reduction in the G3BP1^PS149^ level compared to the CK2α^WT^-expressing neurons (**Fig. 3A, C-D**). These results are further validated by immunoblotting of whole neuronal lysates, which confirm significantly reduced G3BP1^PS149^ levels across all OCNDS-associated CK2α mutant-expressing neurons compared to CK2α^WT^- or GFP-expressing controls (**Fig. S3A-B**). Together, these data indicate that OCNDS-associated CK2α mutations impair G3BP1^PS149^ phosphorylation in a mutation- and compartment-specific manner.

**Figure 3:**
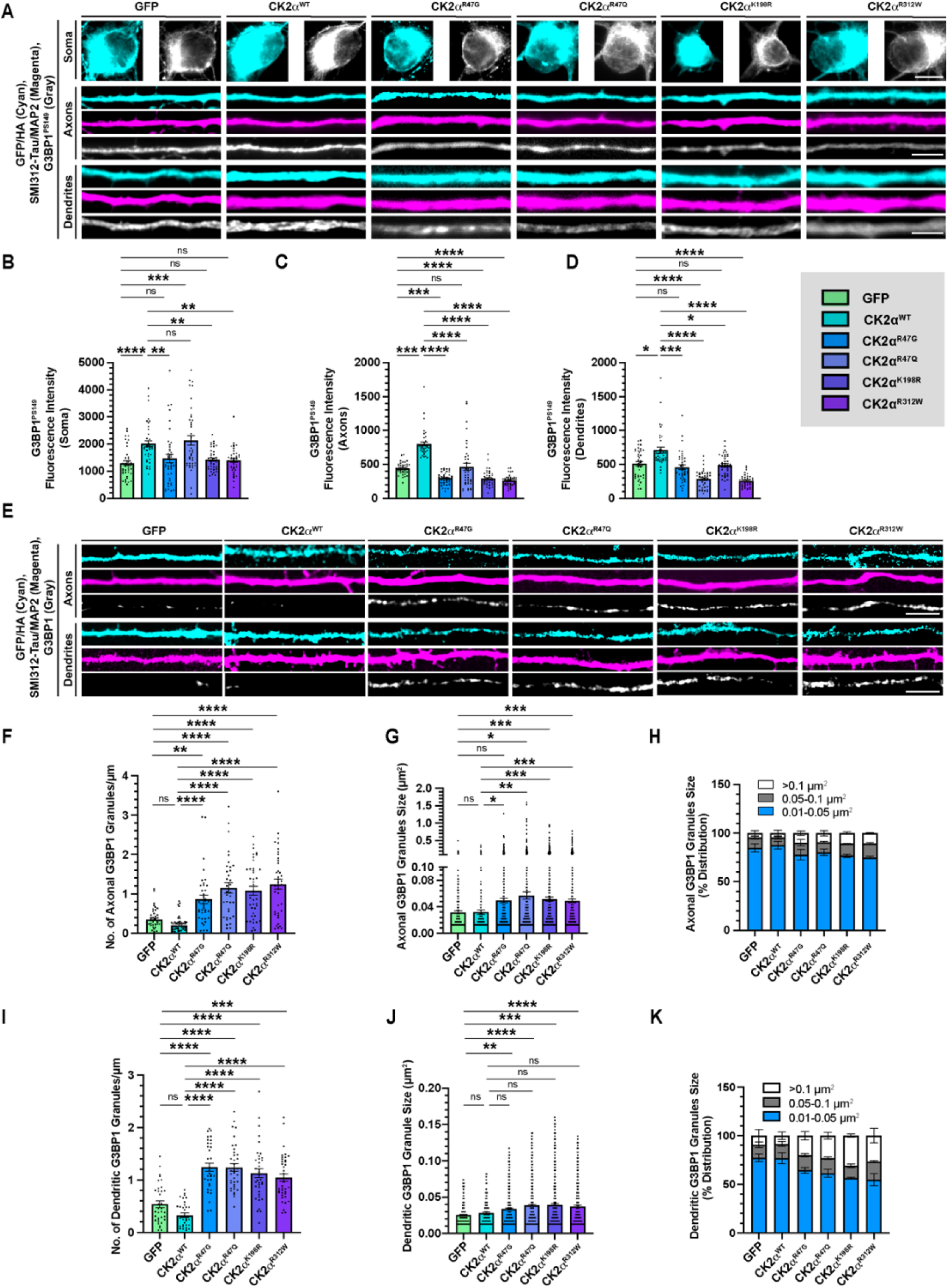
OCNDS-associated CK2α mutants reduce phosphorylated G3BP1^S149^ levels and increase the number and size of axonal and dendritic G3BP1 granules in developing rat cortical neurons. ***A,*** Representative epifluorescent images of soma (top), axons (middle) and dendrites (bottom) of embryonic rat cortical neurons expressing either GFP, CK2α^WT^ or CK2α mutants cultured for 11 *DIV* and immunostained for MAP2, SMI312/Tau, G3BP1^PS149^, and GFP or HA are shown (Scale bar, 10 µm for soma and 5 µm for axons and dendrites). ***B-D,*** Quantifications of phosphorylated G3BP1^S149^ levels in the soma **(B)**, axons **(C)**, and dendrites **(D)** are shown as mean ± SEM. n ≥ 40 neurons across three biological replicates, *p ≤ 0.05, **p ≤ 0.01, ***p ≤ 0.001, ****p ≤ 0.0001 by Kruskal-Wallis test with Dunn’s multiple comparisons. ***E,*** Representative confocal images of axons (top) and dendrites (bottom) of embryonic rat cortical neurons expressing either GFP, CK2α^WT^ or CK2α mutants cultured for 11 *DIV* and immunostained for SMI312/Tau, G3BP1, and GFP or HA are shown. (Scale bar, 5 µm). ***F-H,*** Quantifications of the number of axonal G3BP1 granules/μm **(F)**; n ≥ 36 axons, average granule size **(G)**; n ≥ 174 granules, and percentage distribution of granules as per size **(H)** are shown as mean ± SEM across three biological replicates; *p ≤ 0.05, **p ≤ 0.01, ***p ≤ 0.001, ****p ≤ 0.0001 by Kruskal-Wallis test with Dunn’s multiple comparisons. ***I-K,*** Quantifications of the number of dendritic G3BP1 granules/μm **(I)**; n ≥ 35 dendrites, average granule size **(J)**; n ≥ 186 granules, and percentage distribution of granules as per size **(K)** are shown as mean ± SEM; **p ≤ 0.01, ***p ≤ 0.001, ****p ≤ 0.0001 by Kruskal-Wallis test with Dunn’s multiple comparisons.

Because phosphorylation of G3BP1 at S149 inhibits its propensity for LLPS by increasing the threshold concentration of RNA required to trigger G3BP1 condensation(*46*), we next examined whether the reduced G3BP1^PS149^ levels observed in OCNDS-associated CK2α mutants are associated with changes in SG formation across neuronal compartments. To this end, we assessed the core SG proteins G3BP1 and G3BP2 in neurons expressing CK2α^WT^ or OCNDS-associated CK2α mutants. Our results show an increase in both the density (**Fig. 3E-F, I**) and size (**Fig. 3E, G, J**) of G3BP1 granules in axons and dendrites relative to neurons overexpressing CK2α^WT^. Analysis of granule size distributions (**Fig. 3H, K**) further indicates a shift toward larger granules in CK2α mutant conditions. Similar alterations are also observed for G3BP2 granules (**Fig. S3C-I**). Together, these data suggest that OCNDS-associated CK2α mutations reduce G3BP1^S149^ phosphorylation and promote the formation of larger and more abundant SGs in axons and dendrites.

### Reduced G3BP1^S149^ phosphorylation and increased SG accumulation in OCNDS mouse and human iPSC-derived neuron models

Since we observed an overall reduction in G3BP1^PS149^ levels in cultured developing rat cortical neurons expressing the CK2α mutants *in vitro*, we turned to an *in vivo* OCNDS mouse model carrying the K198R mutation in the *CSNK2A1* gene (*CSNK2A1*^K198R/+^), a knock-in model previously characterized by the Rebholz laboratory(*23*). Quantitative immunostaining revealed that axons and dendrites in the cortex of the *CSNK2A1*^K198R/+^ mouse model show significantly reduced G3BP1^PS149^ levels as compared to WT (**Fig. 4A-C**).

**Figure 4:**
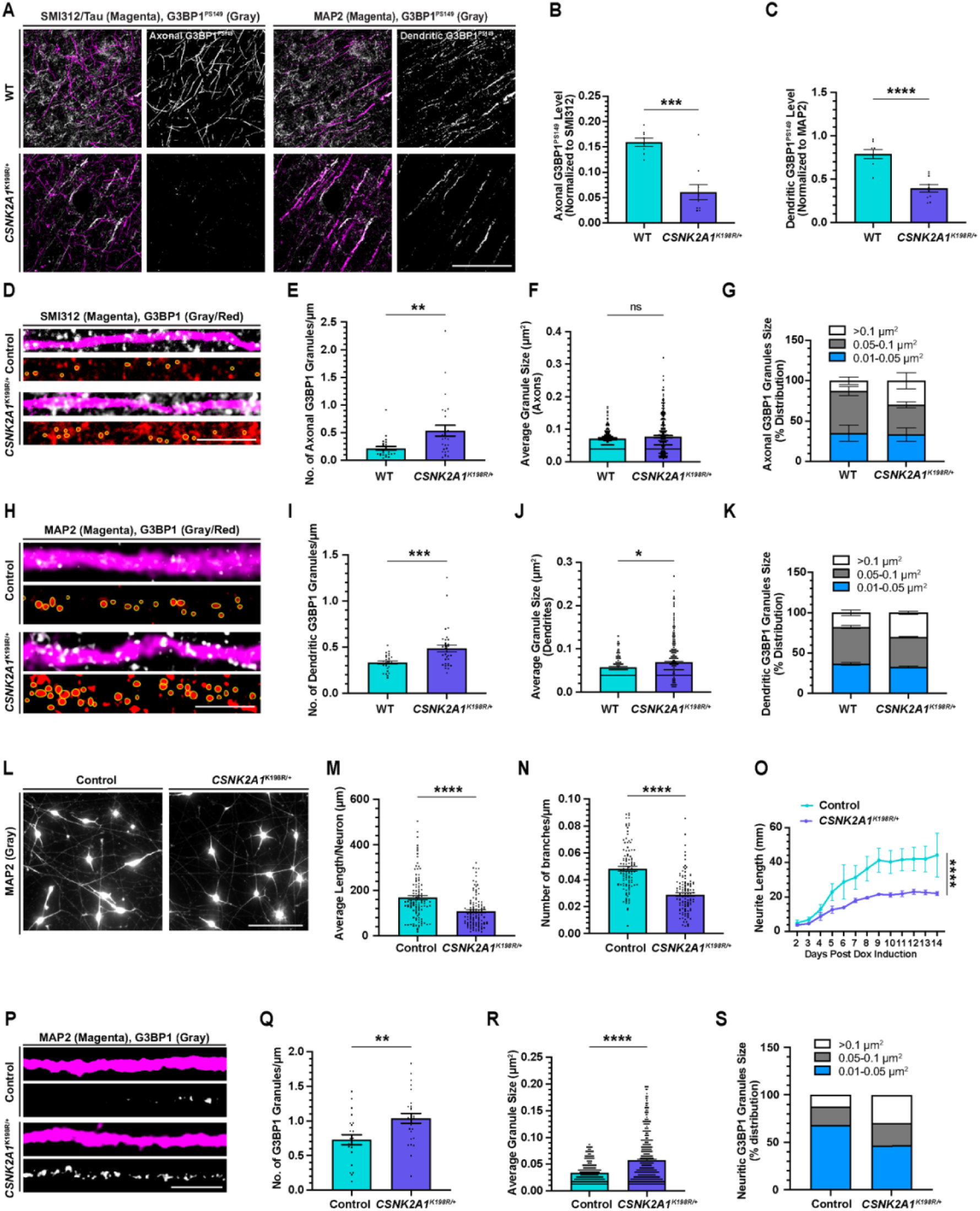
Mouse and human OCNDS models exhibit increased G3BP1 granules and reduced neurite growth. ***A,*** Representative confocal images of axons (left) and dendrites (right) of cortices from coronal sections of adult *CSNK2A1*^K198R/+^ or WT mouse brain immunostained for SMI312/Tau, MAP2, and phosphorylated G3BP1^PS149^ are shown (Scale bar, 50 µm). ***B-C,*** Quantifications of G3BP1^PS149^ levels colocalizing with and normalized to SMI312/Tau for axons **(B)** and MAP2 for dendrites **(C)** are shown as mean ± SEM. n ≥ 8 images from 5 animals of each genotype; ***p ≤ 0.001 by Mann-Whitney for axons, ****p ≤ 0.0001 by Unpaired t-test for dendrites. ***D,*** Representative confocal images of axons of cortices from adult *CSNK2A1*^K198R/+^ or WT mouse brain immunostained for SMI312/Tau and G3BP1 are shown (Scale bar, 5 µm). ***E-G,*** Quantifications of the number of axonal G3BP1 granules/μm **(E)**; n ≥ 28 axons, average granule size **(F)**; n ≥ 121 granules, and percentage distribution of granules as per size **(G)** are shown as mean ± SEM across 5 animals of each genotype; **p ≤ 0.01 **(E)** by Mann-Whitney, and ns **(F)**. ***H,*** Representative confocal images of dendrites of cortices from adult *CSNK2A1*^K198R/+^ or WT mouse brain immunostained for MAP2 and G3BP1 are shown (Scale bar, 5 µm). ***I-K,*** Quantifications of the number of dendritic G3BP1 granules/μm **(I)**; n ≥ 27 dendrites, average granule size **(J)**; n ≥ 224 granules, and percentage distribution of granules as per size **(K)** are shown as mean ± SEM across 5 animals of each genotype; ***p ≤ 0.001 **(I)** and *p ≤ 0.05 **(J)** by Mann-Whitney. ***L,*** Representative epifluorescent images of *CSNK2A1*^K198R/+^ and control iNeurons cultured for 7 *DIV and* immunostained for MAP2 are shown (Scale bar, 100 µm). ***M-N,*** Quantification of average neurite length **(M)** and branching complexity of neurites **(N)** using WIS Neuromath in *CSNK2A1*^K198R/+^ iNeurons are shown as mean ± SEM. n ≥ 111 neurons across three biological replicates; ****p ≤ 0.0001 by Mann-Whitney test. ***O,*** Quantification of average neurite length per square mm of culture well over time taken on Incucyte is shown as mean ± SEM across three biological replicates; ****p ≤ 0.0001 for 9-11 and 13 days post Dox induction by Two-way ANOVA with Tukey’s multiple comparisons. ***P,*** Representative confocal images of neurites of *CSNK2A1*^K198R/+^ and control iNeurons cultured for 7 *DIV and* immunostained for MAP2 and G3BP1 are shown (Scale bar, 5 µm). ***Q-S,*** Quantifications of the number of neuritic G3BP1 granules/μm **(Q)**, n ≥ 23 neurites across three biological replicates, average granule size **(R)** n ≥ 325 granules across three biological replicates, and percentage distribution of granules as per size **(S)** are shown as mean ± SEM across three biological replicates; **p ≤ 0.01, ****p ≤ 0.0001 by Unpaired t-test.

We next examined whether this reduction in G3BP1^PS149^ levels affects G3BP1 granule accumulation in the *CSNK2A1^K198R/+^* cortex. In axons, G3BP1 granule number is significantly increased without a corresponding change in size (**Fig. 4D-G)**, whereas in dendrites, both G3BP1 granule number and size are increased compared to WT (**Fig. 4H-K**). Consistent with these findings, overall G3BP1 granule intensity is elevated in both compartments (**Fig. S4A-C)**. Together, these data demonstrate that reduced G3BP1^PS149^ phosphorylation in the *CSNK2A1^K198R/+^* cortex is accompanied by a compartment-specific increase in G3BP1 granule accumulation, with axons showing increased granule number and dendrites exhibiting both increased number and size *in vivo*.

To assess whether these phenotypes are conserved in humans, we analyzed human induced pluripotent stem cell (hiPSC)-derived iNeurons from an individual with OCNDS carrying the K198R mutation (*CSNK2A1^K198R/+^*) and a control hiPSC line. Neuromath analysis revealed reduced neurite length and branching complexity in *CSNK2A1^K198R/+^* iNeurons compared to the controls (**Fig. 4L-N**), and live-cell imaging using the Incucyte system further demonstrated a reduced neurite growth rate throughout neuronal maturation (**Fig. 4O**). Consistent with our rodent findings, *CSNK2A1^K198R/+^* iNeurons also exhibit increased G3BP1 granule number and size (**Fig. 4P-S**), accompanied by overall higher G3BP1 granule intensity compared to controls (**Fig. S4D-E**). Collectively, these findings establish that OCNDS-associated CK2α dysfunction reduces G3BP1^PS149^ phosphorylation and drives compartment-specific SG accumulation across *in vitro* and *in vivo* rodent models. Notably, the conservation of these phenotypes in patient-derived human neurons, alongside impaired neurite growth, underscores their relevance to the human disorder condition.

### OCNDS-associated CK2α mutants alter G3BP1-mRNA interaction in a compartment-specific manner

As SGs are ribonucleoprotein complexes, we further assessed which axonal mRNAs interact with G3BP1. For this, we performed G3BP1 RNA immunoprecipitation (RIP) followed by bulk RNA sequencing from the axonal compartments of the embryonic rat cortical neurons cultured on porous inserts (**Fig. 5A, Table S1**). RNA-seq analysis revealed that 4290 RNAs are significantly enriched in the G3BP1 immunoprecipitate compared to control IP (**Fig. 5B**). Gene ontology analysis of the G3BP1-interacting RNAs showed strong enrichment for pathways involved in regulation of neuron projection development and intracellular protein transport (**Fig. 5C**), while ClinVar analysis of these mRNAs showed strong enrichment of pathways associated with intellectual disability, neurodevelopmental delay, and neurodevelopmental disorder (**Fig. 5D**). Together, these findings suggest that axonal G3BP1 granules associate with transcripts involved in neuronal growth and maturation.

**Figure 5:**
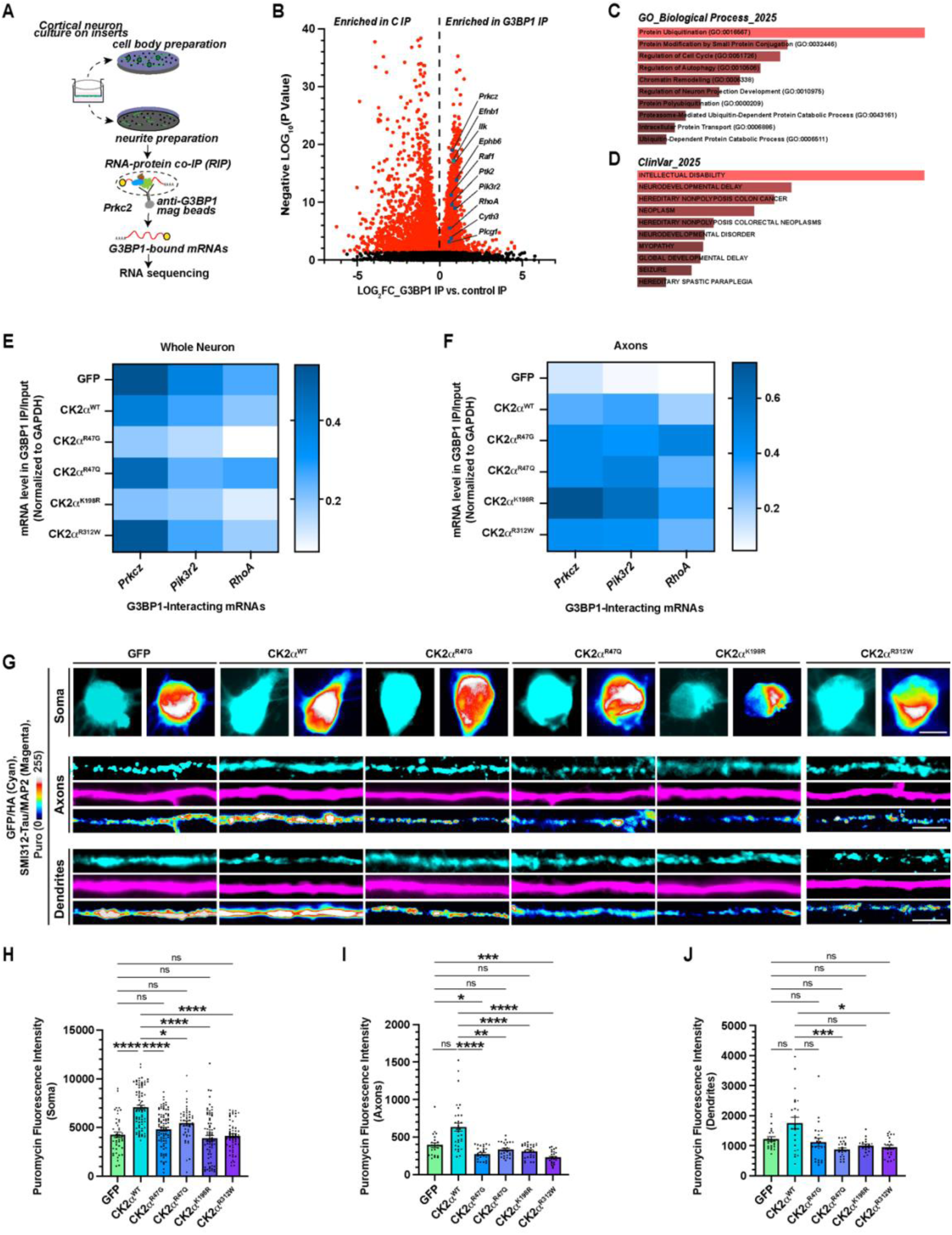
OCNDS-associated CK2α mutants alter interactions of specific mRNAs with G3BP1 granules in a compartment-specific manner and inhibit protein synthesis in developing rat cortical neurons. ***A,*** Schematic showing the experimental design for axonal G3BP1-RNA co-IP followed by bulk RNA sequencing. ***B,*** Volcano plot shows mRNAs enriched in the G3BP1 immunoprecipitate. Significantly enriched mRNAs are highlighted in red, and the mRNAs selected for downstream analysis are marked. ***C-D,*** Gene Ontology (GO) Biological Process enrichment analysis (**C**), and ClinVar disease enrichment analysis (**D**) of mRNAs significantly associated with G3BP1 are shown. ***E-F,*** Heat maps show the quantifications of mRNA levels in the indicated expression conditions from whole neurons **(E)** and axons **(F)** following G3BP1 immunoprecipitation and RT-ddPCR analysis are shown as the mean across three biological replicates. ***G,*** Representative epifluorescent images of soma (top), axons (middle) and dendrites (bottom) of embryonic rat cortical neurons expressing either GFP, CK2α^WT^ or CK2α mutants cultured for 11 *DIV*, treated with puromycin, and immunostained for puromycin, SMI312/Tau or MAP2 and GFP or HA are shown (Scale bar, 10 µm for soma and 5 µm for axons and dendrites). ***H-J,*** Quantifications of puromycin levels in the soma **(H)**, axons **(I)**, and dendrites **(J)** are shown as mean ± SEM. n ≥ 43 for soma, n ≥ 72 for axons, and n ≥ 62 dendrites across three biological replicates; *p ≤ 0.05, **p ≤ 0.01, ***p ≤ 0.001, ****p ≤ 0.0001 by Kruskal-Wallis test with Dunn’s multiple comparisons.

We further focused on ten mRNAs enriched in the G3BP1 immunoprecipitate (*Efnb1, Prkcz, Pik3r2, Ilk, Ephb6, Raf1, Ptk2, Cyth3, Plcg1, RhoA*), each previously implicated in neurodevelopment(*48–60*). To validate the interaction of G3BP1 with these ten mRNAs, and to assess the effect of the CK2α mutants on this interaction, we expressed GFP, CK2α^WT^, CK2α^R47G^, CK2α^R47Q^, CK2α^K198R^, or CK2α^R312W^ in cultured rat cortical neurons and performed G3BP1 RIP at 9 *DIV*, validated by immunoblotting (**Fig. S5A**). Reverse transcriptase-coupled droplet digital PCR (RT-ddPCR) analysis of whole neurons revealed that CK2α^R47G^- and CK2α^K198R^-expressing neurons consistently reduce G3BP1 interaction with all ten target mRNAs tested (**Fig. 5E, Fig. S5B**). Given that we had observed increased G3BP1 granule accumulation in neuronal compartments expressing CK2α mutants, this result was unexpected and prompted us to examine the axonal compartment specifically (**Fig. 5A, Fig. S5C)**. As the axonal material is limited, we focused on three mRNAs (*Prkcz*, *Pik3r2*, and *RhoA*) with the strongest prior evidence for axonal mRNA localization and local translation in the context of axon growth and cytoskeletal regulation(*57, 59, 61*). We find that expression of CK2α^WT^ or any mutant increase G3BP1 interaction with all three transcripts in axons (**Fig. 5F**). Notably, CK2α^K198R^ further enhance the interaction of *Prkcz* and *Pik3r2* while CK2α^R47G^ selectively enhanced the interaction of *RhoA*, relative to CK2α^WT^ (**Fig. 5F**). We noticed that although whole-neuron transcript levels show modest mutation-dependent changes, axonal fractions reveal stronger divergence among CK2α variants, particularly for *Pik3r2* and *RhoA* (**Fig. S5D-E**), as expression of CK2α^R47G^ and CK2α^R312W^ reduce axonal *Pik3r2* and *RhoA* levels. Together, these findings demonstrate that OCNDS-associated CK2α mutations differentially regulate the axonal abundance and G3BP1 association of neurodevelopmentally relevant mRNAs in a mutation- and transcript-specific manner. This suggests that dysregulated axonal mRNA-granule dynamics may contribute to the neuronal growth deficits observed in OCNDS.

### OCNDS-associated CK2α mutants suppress local protein synthesis

SGs are known to suppress mRNA translation(*45, 62, 63*), and we have previously shown that disassembling G3BP1 granules increases local protein synthesis in the axons of developing rat cortical neurons(*27, 29*). Given that CK2α mutations increase G3BP1 granule accumulation, we next asked whether they affect neuronal protein synthesis. To this end, we assessed nascent protein synthesis using a puromycinylation assay in 11 *DIV* cultured embryonic rat cortical neurons expressing GFP, CK2α^WT^, or the OCNDS-associated CK2α^R47G^, CK2α^R47Q^, CK2α^K198R^, or CK2α^R312W^ mutants (**Fig. 5G, Fig. S5F-H**). CK2α^WT^ overexpression significantly increased translation in the soma compared to GFP, though this effect did not extend to axons or dendrites (**Fig. 5G-J)**. In contrast, all CK2α mutant-expressing neurons exhibit reduced puromycin signal across all compartments, soma, axons, and dendrites, compared to CK2α^WT^-expressing neurons (**Fig. 5G-J**), indicating that OCNDS-associated CK2α mutations broadly suppress local protein synthesis in developing rat cortical neurons.

### Knockdown of *G3bp1* restores protein synthesis and neuronal growth defects in neurons expressing OCNDS-associated CK2α mutants

Having established that OCNDS-associated CK2α mutations increase G3BP1 granule accumulation and suppress protein synthesis (**Figs. 3 and 5**), we next asked whether reducing G3BP1 levels could rescue translational deficits in these neurons. Neurons expressing GFP, CK2α^WT^, or CK2α mutants were treated with siRNA targeting *G3bp1*, and knockdown efficiency was validated by RT-ddPCR and quantitative immunofluorescence (**Fig. 6A, Fig. S6A-C**).

**Figure 6:**
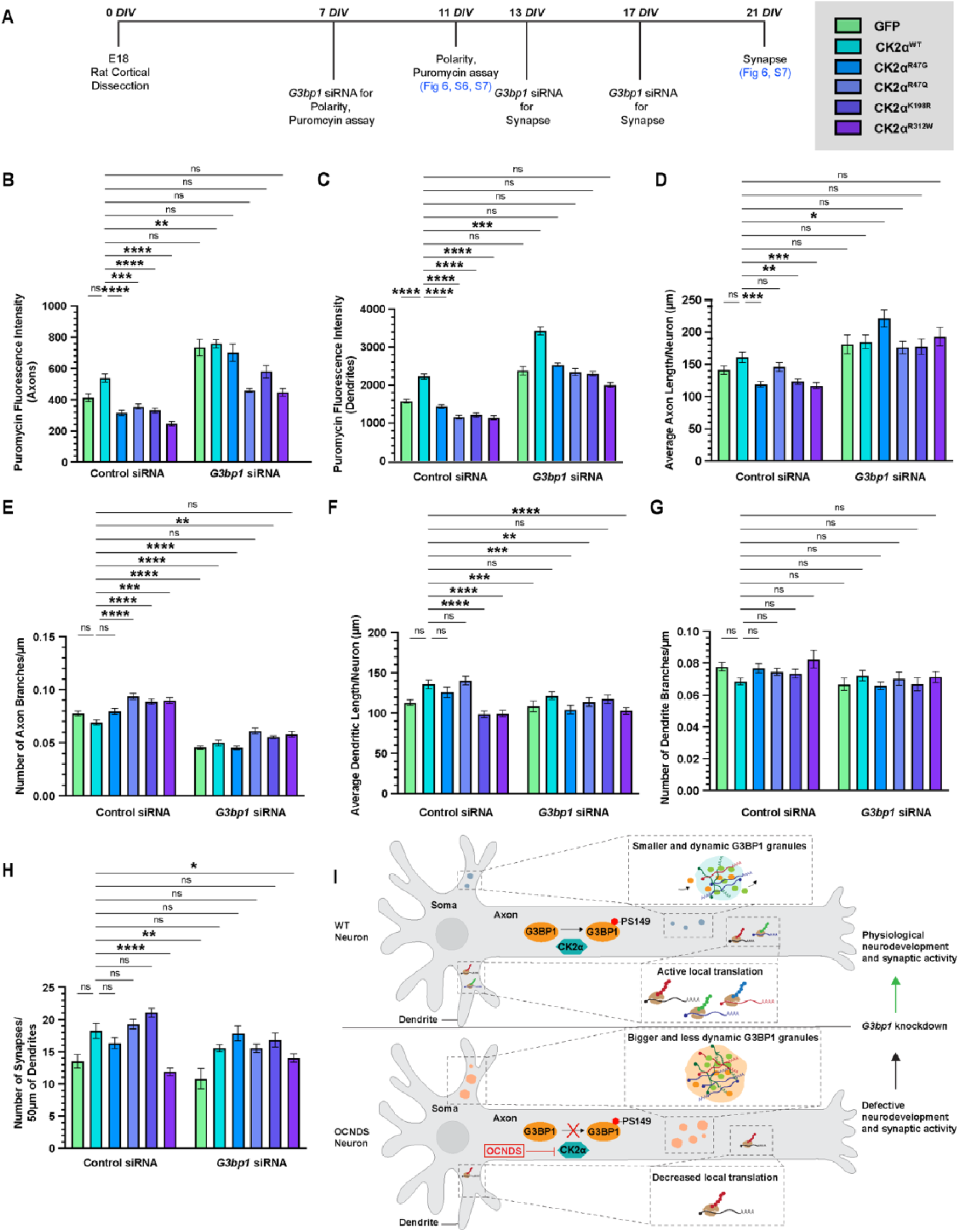
Knocking down G3bp1 rescues defects in protein synthesis, neurite complexity, and synaptogenesis. ***A,*** Timeline for *G3bp1* siRNA-treated E18 rat cortical neurons expressing either GFP, CK2α^WT^ or CK2α mutants and cultured for various timepoints and experiments is shown. ***B-C,*** Quantifications of puromycin levels in the axons **(B)** and dendrites **(C)** following treatment with either control or *G3bp1* siRNA are shown as mean ± SEM. n ≥ 51 axons and n ≥ 62 dendrites across three biological replicates; *p ≤ 0.05, **p ≤ 0.01, ***p ≤ 0.001, ****p ≤ 0.0001 by Kruskal-Wallis test with Dunn’s multiple comparisons. ***D-G,*** Quantifications of the average axonal length **(D),** axonal branching complexity **(E),** average dendritic length **(F),** and dendritic branching complexity **(G)** of axons and dendrites of embryonic rat cortical neurons expressing either GFP, CK2α^WT,^ or CK2α mutants cultured for 11 *DIV,* treated with *control or G3bp1* siRNA on 7 *DIV* for 4 *DIV* are shown as mean ± SEM. n ≥ 72 images across three biological replicates; *p ≤ 0.05, **p ≤ 0.01, ***p ≤ 0.001, ****p ≤ 0.0001 by Kruskal Wallis test with Dunn’s multiple comparisons. ***H,*** Quantification of the number of synapses in embryonic rat cortical neurons expressing either GFP, CK2α^WT^, or CK2α mutants cultured for 21 *DIV* is shown as mean ± SEM. n ≥ 30 neurites; *p ≤ 0.05, **p ≤ 0.01, ****p ≤ 0.0001 by Kruskal-Wallis test with Dunn’s multiple comparisons. ***I,*** Working model depicting the OCNDS-associated CK2α mutants with reduced kinase activity, which cause increased axonal G3BP1 granules (reduced dynamicity) and decreased axonal and dendritic local translation, leading to defects in neurodevelopment and synaptic activity.

Knockdown of *G3bp1* robustly restores puromycin levels in both axons and dendrites across all CK2α mutant conditions, confirming rescue of local protein synthesis (**Fig. 6B-C, Fig. S6D-E**). Notably, *G3bp1* knockdown also elevates translation in GFP- and CK2α^WT^-expressing neurons, suggesting that G3BP1 constitutively suppresses baseline protein synthesis even under physiological conditions. Consistent with this translational recovery, *G3bp1* knockdown increases axonal length and reduces branching density across GFP-, CK2α^WT^-, or CK2α mutant-expressing neurons (**Fig. 6D-E, Fig. S7**). Interestingly, we observe a bidirectional normalization for dendritic morphology. *G3bp1* knockdown abolishes the increased dendritic length seen in CK2α^WT^-, CK2α^R47G^-, and CK2α^R47Q^-expressing neurons, while simultaneously rescuing the dendritic length reductions observed in CK2α^K198R^ and CK2α^R312W^-expressing neurons (**Fig. 6F-G, Fig. S7**). Similarly, *G3bp1* knockdown reduces the excess synapses in CK2α^K198R^-expressing neurons and rescues the deficit in CK2α^R312W^-expressing neurons (**Fig. 6H, Fig. S7**). Together, these findings demonstrate that G3BP1-mediated translational regulation is a central bidirectional driver of the synaptic and morphological phenotypes observed across OCNDS-associated CK2α mutations, positioning G3BP1 as a key downstream effector of CK2α activity in developing neurons (**Fig. 6I**).

## DISCUSSION

Here, we report that OCNDS-associated mutations in *CSNK2A1* converge on a common cellular mechanism, the dysregulation of G3BP1-dependent SG dynamics and compartment-specific suppression of local mRNA translation in developing neurons. We demonstrate that OCNDS-associated CK2α variants, CK2α^R47G^, CK2α^R47Q^, CK2α^K198R^, and CK2α^R312W^, reduce G3BP1 phosphorylation at serine 149 (G3BP1^PS149^). This promotes the formation of larger, more persistent, and less dynamic SGs (**Figs. 1-4**). These granules sequester neurodevelopmentally relevant mRNAs and suppress local protein synthesis across all neuronal compartments (**Fig. 5**). Knockdown of *G3bp1* rescues both translational deficits and the morphological and synaptic phenotypes across all mutants tested (**Fig. 6**). These findings establish G3BP1-mediated translational regulation as a central effector of CK2α function in developing neurons.

OCNDS-associated CK2α mutants increase SG number and size and reduce SG dynamicity, even in the absence of exogenous stress. This extends prior work showing that pharmacological inhibition of CK2α enhances SG formation in melanoma cells(*40*). CK2α phosphorylates G3BP1 at S149 to raise the RNA concentration threshold for phase separation and promote granule disassembly(*20, 28, 46*). All four OCNDS-associated CK2α mutants tested significantly reduced G3BP1^PS149^ levels in axons and dendrites, leading to more abundant granules in both compartments. The reduction in G3BP1-G3BP2 interaction observed in co-immunoprecipitation experiments, particularly in CK2α^K198R^ and CK2α^R312W^ mutants (**Fig. 1E-G**), further suggests that the SG core is possibly defective in the cells expressing these mutants(*44–46*). Together, these data position G3BP1^PS149^ as a critical regulator downstream of CK2α, tuning the assembly-disassembly equilibrium of neuronal granules. At the morphological level, CK2α^K198R^ and CK2α^R312W^, which broadly reduced G3BP1^PS149^, showed the most consistent deficits in axonal length and dendritic complexity. In contrast, CK2α^R47Q^ preserved somatic G3BP1^PS149^ and increased dendritic growth, suggesting that residual kinase activity partially spares dendritic development (**Figs. 2-3**). CK2α^K198R^ caused an increased number of synapses, while CK2α^R47G^ and CK2α^R312W^ caused synaptic loss (**Fig. 2**). This bidirectionality reflects engagement of distinct downstream signaling nodes and is consistent with the clinical heterogeneity observed across OCNDS alleles(*20, 24, 47*)

CK2α^K198R^ expression drives not only synaptic overabundance but also increased and irregular spontaneous network activity. Elevated Fano Factors at 21 *DIV* indicate heightened firing variability (**Fig. 2I-K**). These electrophysiological features are reminiscent of hyperexcitable network states in ASD and intellectual disability(*64–67*), and are consistent with the seizure phenotypes reported in OCNDS patients(*6, 7, 23, 68, 69*). Overexpression of CK2α^WT^ transiently elevates spike rates at 14 *DIV* but normalizes by 21 *DIV*. CK2α^K198R^-expressing neurons fail to return to baseline firing by this time point. This suggests that physiological CK2α activity is required to calibrate network maturation within a critical developmental window. Loss of this activity may lock neurons in a hyperexcitable immature state through the combined effects of excess synapses and impaired translational homeostasis.

Expression of all OCNDS-associated CK2α mutants broadly suppresses local protein synthesis across soma, axons, and dendrites (**Fig. 5G-J**). Local translation in axons and dendrites is critical for the autonomous regulation of neuronal growth, synaptic plasticity, and injury responses(*27, 29, 70–72*). This is consistent with our previous demonstration that CK2α-mediated G3BP1 granule disassembly is a key mechanism for the release of axonal mRNAs for translation following sciatic nerve injury(*28*). Axonal G3BP1-associated transcripts are enriched for gene ontology terms related to neuron projection development and for ClinVar associations with intellectual disability and neurodevelopmental delay (**Fig. 5B-D**). Individual mutants selectively trap specific transcripts in granules. For example, expression of CK2α^K198R^ enhances G3BP1 interaction with *Prkcz* and *Pik3r2*, while expression of CK2α^R47G^ selectively enhances interaction with *RhoA* in axons (**Fig. 5F**). This provides a mechanistic basis for the distinct morphological phenotypes observed across mutants. Moreover, individual OCNDS-associated CK2α mutants differentially alter the axonal abundance and G3BP1 association of specific mRNA populations (**Fig. S5A-E**). This suggests that each mutant reshapes the local axonal translatome in a distinct manner. How individual mutants restructure granule-associated transcriptomes and proteomes, and what additional contributions arise from other CK2α substrates, remain important questions for future studies.

Reduced G3BP1^PS149^ and increased granule accumulation are conserved across three independent model systems, including rat cortical neurons *in vitro*, the *CSNK2A1^K198R/+^* knock-in mouse cortex *in vivo*, and *CSNK2A1^K198R/+^* iNeurons (**Fig. 4**), supporting the relevance of these findings across rodents and humans. Notably, knockdown of *G3bp1* restored puromycin incorporation in axons and dendrites across all mutants, elevating the baseline translation in GFP- and CK2α^WT^-expressing neurons, confirming that G3BP1 suppresses local protein synthesis even under physiological conditions(*^41,42, 46^*). *G3bp1* knockdown bidirectionally normalized dendritic morphology, abolishing excess growth in CK2α^R47G^/CK2α^R47Q^-expressing neurons and rescuing growth inhibition in CK2α^K198R^/CK2α^R312W^-expressing neurons (**Fig. 6F-G**). It also rescued synaptic excess in CK2α^K198R^-expressing neurons and synaptic deficit in CK2α^R312W^-expressing neurons (**Fig. 6H**). These findings indicate that synaptic and morphological phenotypes are downstream of translational dysregulation, rather than direct consequences of impaired CK2α substrate phosphorylation at the synapse.

Although CK2α has nearly five hundred known substrates(*20, 47*), the data presented here indicate that its control of G3BP1 phase separation is a functionally dominant axis in developing neurons. This raises the possibility that other kinases implicated in neurodevelopmental disorders may similarly regulate granule dynamics(*73, 74*), and that aberrant phase separation may be a shared convergent mechanism across a wider class of such conditions(*75–78*). The robust rescue of translational, morphological, and synaptic phenotypes by *G3bp1* knockdown identifies the CK2α–G3BP1 axis as a candidate therapeutic target. In summary, OCNDS-associated CK2α mutations converge on G3BP1-mediated translational repression, thereby impairing neuronal morphogenesis and synaptic development. More broadly, these results highlight the importance of local translational control in neurodevelopment and suggest that dysregulation of RNA granule homeostasis may be a common pathogenic thread among intellectual disability syndromes.

## ONLINE METHODS

### For *in vitro* and *in vivo* animal studies

The Institutional Animal Care and Use Committee of Rutgers University and Université de Paris approved all animal procedures. Timed-pregnant Sprague Dawley rats and adult WT and *CSNK2A1^K198R/+^*heterozygous littermates from C57BL/6J-Csnk2a1em2Lutzy/Mmjax mice (MMRRC stock #67399, The Jackson Laboratory) were used for all experiments.

### For studies involving stem cells

All human induced pluripotent stem cell (hiPSC) experiments were conducted following prior approval from the University of Michigan Human Pluripotent Stem Cell Research Oversight (HIPSCRO) Committee. CC1A control cell line was a gift from Dr. Andrew Tidball (University of Michigan). CSNK2A1 593A>G iPSC cell line (SV0002578) was acquired through the Simons Foundation Autism Research Initiative (SFARI) cell repository.

### hiPSC culture and iNeuron generation

All cells were maintained in mTeSR™Plus (STEMCELL Tech; 100-0274) at 37 °C and 5% CO_2_. Cells were passaged regularly by washing with DPBS (Gibco™; 14190144) and incubated in 0.8 mM EDTA (Lonza; 51201) for 4 min. SV0002578 cells were transfected with the UCMCLYBL-Ngn1/2 plasmid, a gift from Dr. Michael Ward (National Institute of Neurological Disorders and Stroke) and the 2 CLYBL targeting TALEN plasmids (pZT-C13-R1 and pZT-C13-L1, gifts from Jizhong Zou (National Heart Lung and Blood Institute). Addgene.org: 52638 and 52637, respectively) using Lipofectamine stem (Invitrogen; STEM00003). Transfected clones were single cell selected using expressed mCherry and insertion number was validated via PCR. CC1A line had been transfected previously(*79, 80*). The SV0078N1 clone was selected for this study based on PCR results and morphology. To form neurons, cells were passaged to single cells using StemPro™ Accutase™ (Gibco™; A11105-01) for 7 min at 37 °C. Cells were then replated in mTeSR^™^ Plus containing 10 µM Rho Kinase inhibitor (Y-27632; Cayman Chemical; 10005583) at a density of 5 x 10^5^ cells/well of a 6-well plate. The next two days, the medium was removed and replaced with 1mM Doxycycline (DOX) (Cayman Chemical; 14422). On the third day, the cells were single-cell passaged again at a density of 4000 or 1000 cells per well for 12-well and 96-well plate, respectively into DOX-containing medium. The following day, the medium was changed to 3N+A (*81*) + 1 mM DOX. This was continued every other day for the remainder of the experiment.

### For cell lines and primary cultures

COS-7 and HEK293T cells were maintained in DMEM (HyClone™; 16750-074) supplemented with 10% fetal bovine albumin (FBS) (Sigma; F2442) and 1% penicillin/streptomycin (P/S) (Gibco™; 15140122). Cells were passaged using 0.25% Trypsin (Gibco™; 15050057) at 70-90% confluency.

For cortical neuron cultures, E18 rat cortices were dissected in Hank’s Balanced Salt Solution (HBSS) (Life Technologies; 45000-462) and dissociated using Trypsin as described previously(*27, 82*). Briefly, cortices were minced and incubated in pre-warmed 0.25% trypsin at 37 °C for 5 min. After two washes in HBSS at 37 °C for 5 min each, tissues were triturated in Neurobasal medium (Life Technologies; 21103049) with 1x L-Glutamine (Life Technologies; 25-030-081) and 1x B27 supplement (Life Technologies; 17504-044), and post-nucleofection, plated on acid-rinsed coverslips (Chemglass Life Sciences; CLS-1760-012) for immunostaining, polyethylene-tetrathalate (PET) membrane inserts (1 μm pores; Corning; 62406-171) for isolation of axons from cell bodies, or tissue culture-treated plates for isolation of protein or RNA-immunoprecipitation (RIP), all pretreated with 100 μg/mL poly-L-lysine (Sigma Aldrich; P1274). Approximately 2 h after plating, the medium was changed to fresh Neurobasal supplemented with 1x L-Glutamine, 1x B27, and 1 μg/mL laminin (Life Technologies; 23017-015). Cells were cultured at 37 °C in 5% CO_2,_ and half the medium was changed every 3 days for 9-21 days *in vitro (DIV)*, depending on the experiment. Axon and cell body preparations were isolated from neurons grown on PET membranes as described previously(*83*) for RIP.

### Generation of OCNDS-Associated CK2α mutant constructs

OCNDS-associated mutations in *CSNK2A1* encoding the catalytic subunit of CK2α were introduced into a C-terminal 3x HA-tagged CK2α expression construct using site-directed mutagenesis. Human CK2α-HA construct was from Addgene (plasmid 27086; RRID: Addgene_27086; originally generated by David Litchfield) and served as the template for all mutagenesis reactions. Site-directed mutagenesis was performed using the Q5 Site-Directed Mutagenesis Kit (New England Biolabs (NEB), E0554S) according to the manufacturer’s instructions. This kit utilizes high-fidelity amplification with Q5 Hot Start High-Fidelity DNA Polymerase, followed by kinase-ligase-DpnI (KLD) treatment to circularize amplified plasmids and remove methylated parental DNA.

### Primer design

Mutagenic primers were designed using the NEBaseChanger primer design tool provided by NEB. Primers were designed to introduce the desired missense mutations within the HA-tagged CK2α construct, while maintaining optimal melting temperature and amplification efficiency. The primer sequences used to generate the CK2α mutants are listed below:

### *CSNK2A1* Mutation Primers

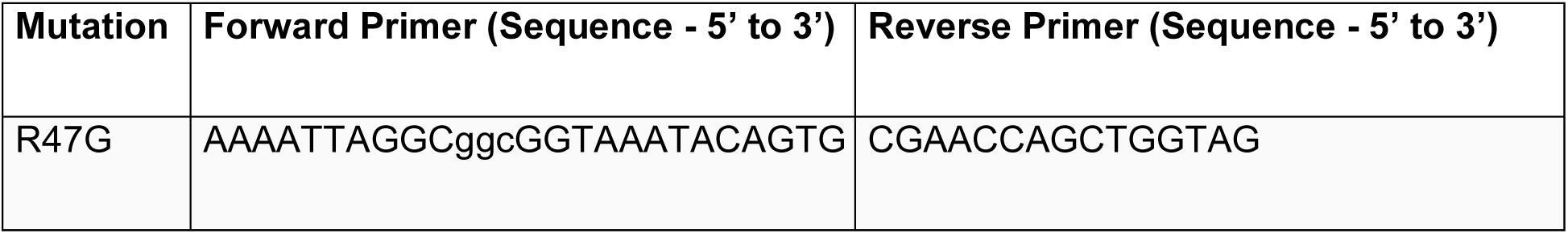
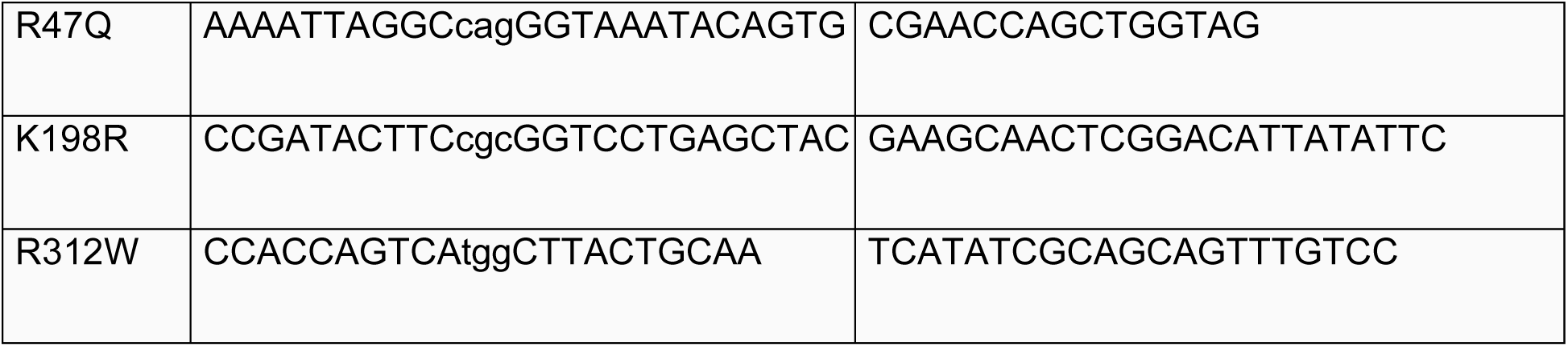

### Site-directed mutagenesis by PCR and KLD treatment

Site-directed mutagenesis was performed using PCR amplification in 25 µL reaction volumes containing 12.5 µL of 2x Q5 Hot Start High-Fidelity Master Mix, 1.25 µL each of forward and reverse mutagenic primers (10 µM), 5 ng of HA-CK2α^WT^ plasmid template DNA, and nuclease-free water. PCR amplification was carried out in a thermocycler under the following conditions: an initial denaturation at 98 °C for 30 s, followed by 25 cycles of denaturation at 98 °C for 10 s, annealing at 60-65 °C for 20 s (depending on primer melting temperature), and extension at 72°C for 2-3 min (30 s per kb of plasmid length), with a final extension at 72 °C for 2 min and a hold at 4°C. Following amplification, the PCR product was subjected to kinase-ligase-DpnI (KLD) treatment to phosphorylate, ligate, and digest the parental template DNA. The KLD reaction mixture consisted of 5 µL PCR product, 1 µL 10x KLD reaction buffer, 1 µL KLD enzyme mix, and 3 µL nuclease-free water, and was incubated at room temperature for 5 min prior to downstream applications.

### Bacterial transformation

Following KLD treatment, 5 µL of the reaction mixture was transformed into chemically competent DH5α *Escherichia coli* (*E. coli*; NEB; C29871). Cells were thawed on ice, mixed with the KLD reaction, incubated on ice for 30 min, then heat-shocked at 42 °C for 30 s, followed by recovery on ice for 5 min. Subsequently, 250 µL of SOC medium was added, and cells were incubated at 37 °C with shaking for 1 h. Transformed bacteria were plated onto LB agar plates containing 100 µg/mL ampicillin and incubated overnight at 37 °C.

### Plasmid isolation and sequence verification

Individual colonies were picked and cultured in 3-5 mL LB medium with ampicillin selection overnight at 37 °C with shaking. Plasmid DNA was isolated using a miniprep plasmid purification kit (ZymoResearch; D4210). The presence of the desired mutations and the absence of unintended mutations were confirmed by Sanger sequencing with primers flanking the mutation. All confirmed constructs were expanded and purified for downstream experiments.

### Cloning of mCherry-tagged CK2α constructs

The coding sequences of CK2α-HA and the mutant CK2α^K198R^-HA were subcloned into a lentiviral expression vector (pLV_mCherry: Addgene plasmid 36084) to generate C-terminal mCherry-tagged constructs. Both insert and vector were digested using the restriction enzymes BamHI and XbaI. The pLV_mCherry vector backbone was linearized by double digestion with BamHI and XbaI. In parallel, CK2α-HA and CK2α^K198R^-HA inserts were excised from their original plasmids using the same restriction enzymes to ensure compatible cohesive ends. Following digestion, vector and insert fragments were purified using a gel extraction kit.

Ligation was performed using T4 DNA Ligase (NEB; M0202) according to the manufacturer’s instructions. Briefly, vector and insert were combined at a molar ratio of approximately 1:3 in the presence of 1x T4 DNA Ligase Reaction Buffer and 400 U T4 DNA Ligase in a total reaction volume of 10-20 µL. The ligation mixture was incubated at 16 °C overnight (or alternatively at room temperature for 10-15 min for rapid ligation). Following ligation, constructs were transformed into chemically competent *E. coli*, and positive clones were selected and verified by restriction digestion and sequencing. This cloning strategy resulted in expression constructs encoding mCherry-tagged CK2α^WT^ and CK2α^K198R^ mutant proteins.

### Transfection

For granularity analysis, 5 x 10^4^ COS-7 cells were plated on acid-rinsed coverslips in 24-well plates in DMEM supplemented with 10% FBS and 1% P/S, and after 3 h, they were transfected with 100 ng HA-tagged CK2α^WT^, CK2α^R47G^, CK2α^R47Q^, CK2α^K198R^, CK2α^R312W^ or GFP construct using the calcium phosphate mammalian transfection kit (Takara Bio; 631312). Approximately 16 h after plating, the medium was changed, and the cells were maintained for an additional 24 h for further use.

For fluorescence recovery after photobleaching (FRAP) experiments, 1.5 x 10^4^ COS-7 cells were plated in live cell dishes (FluoroDish™; FD35-100 in DMEM supplemented with 10% FBS and 1% P/S, and after 3 h, transfected with 100 ng GFP-G3BP1 and 300 ng mCherry-tagged CK2α^WT^, CK2α^K198R^ or mCherry (control) constructs using a calcium phosphate mammalian transfection kit. Approximately 16 h after plating, the medium was changed to phenol-free DMEM medium (Gibco; 21041025), and the cells were maintained for an additional 24 h for further use.

For immunoprecipitation (IP) 0.5 x 10^6^ HEK293T cells were plated in 6-well plates in DMEM supplemented with 10% FBS and 1% P/S, and after 3 h, they were transfected with 0.5 μg plasmid HA-tagged CK2α^WT^, CK2α^R47G^, CK2α^R47Q^, CK2α^K198R^, CK2α^R312W^ or GFP construct using calcium phosphate mammalian transfection kit.

For cortical neuron transfection, 4.5 x 10^6^ cells/transfection were pelleted by centrifugation at 80 x *g* for 5 min and resuspended in ‘nucleofector solution + supplement’ (Rat Neuron Nucleofector kit; Lonza; VPG-1003). 2 μg CK2α or 3 μg GFP plasmids were electroporated into the neurons using an AMAXA Nucleofector apparatus (program Neurons Rat Hippocampal neurons, G-013; Lonza)(*28*).

For siRNA transfection, 125 nM On-target plus-SMART pool siRNA against *G3bp1* mRNAs (Dharmacon; L-101659-02-0005) were used with DharmaFECT 3 reagent (Dharmacon; T-2003-02) and incubated for 96 h (*28*). DharmaFECT 3 reagent with siRNA buffer was used as a control.

### Sodium arsenite stress and recovery

The transfected COS-7 cells were stressed with 0.5 mM sodium arsenite (Sigma Aldrich; S7400) for 30 min, and then the medium was completely changed for stressed cells to study SG assembly and disassembly. The recovery was monitored at 0 min, 60 min, and 120 min.

### Fluorescence Recovery After Photobleaching (FRAP) assay

FRAP was used to assess the dynamicity of SGs. COS-7 cells were co-transfected with plasmids encoding GFP-tagged G3BP1 and mCherry-tagged CK2α^WT^, and CK2α^K198R^ and mCherry as a control using calcium phosphate mammalian transfection kit. The COS-7 cells co-transfected with GFP-tagged G3BP1 and mCherry-tagged CK2α^WT^, and CK2α^K198R^ and mCherry constructs were stressed with 0.5 mM sodium arsenite for 30 min before FRAP acquisition. Cells were maintained at 37° C, 5% CO_2_ during imaging sequences using Zeiss LSM980 Confocal Microscope with Airyscan 2. A 488 nm laser line with a laser power of 100% was used to bleach GFP signal post first image for a duration of 0.5 ms, and its recovery was observed with 63X oil immersion objective. The pinhole was set to 1 Airy unit to ensure full thickness bleaching and acquisition. Regions of interest (ROIs) were marked for moderately sized GFP-positive SGs, specifically in mCherry-transfected cells (small SGs were avoided because they were highly dynamic and would escape the ROIs over time). Prior to photobleaching, co-transfected COS-7 cells were imaged after 2s to acquire baseline fluorescence of the ROIs (0.2% laser power at 488nm for GFP and 2% at 561nm for mCherry). The same excitation and emission parameters were used to assess recovery for GFP signal over 150 s post-bleach with images acquired at 2 s intervals. Fluorescence intensities in the ROIs were calculated by the ZEISS software. To normalize across experiments, the fluorescence intensity at t=0 s post-bleach was subtracted from each image sequence to set the baseline to 0%. The percentage of fluorescence recovery at each time point after photobleaching was then calculated by normalizing relative to the pre-bleach fluorescence intensity (set as 100%).

### IP and immunoblotting

HEK cells were collected 48 h after transfection, washed twice with PBS, and lysed by rotation in IP buffer containing 150 mM NaCl, 10 mM Tris-HCl, 1 mM EDTA, 1 mM EGTA, 1% Triton X-100, 0.5% NP-40, and 1 mM DTT, supplemented with 1x protease inhibitor cocktail (Roche; 4693116001). Lysates were clarified by centrifugation at 10,000 x *g* for 15 min at 4 °C. Cleared supernatants were incubated with rabbit anti-G3BP1 antibody (1 µg; Sigma; G6046) for 2 h at 4°C, followed by immunoprecipitation with Protein A/G magnetic beads (Invitrogen; 88802) for an additional 2 h with rotation. Bead-only samples processed in parallel without primary antibody were used as negative controls. Beads were washed four times with cold IP buffer.

For immunoblotting, whole-cell lysates or immunoprecipitated samples were denatured by boiling in Laemmli sample buffer, resolved by SDS-PAGE, and transferred onto nitrocellulose membranes. Membranes were blocked for 1 h at RT in 5% bovine serum albumin (BSA) prepared in Tris-buffered saline containing 0.1% Tween-20 (TBST), followed by incubation with primary antibodies diluted in 5% BSA overnight at 4 °C with gentle rocking. Primary antibodies used were rabbit anti-G3BP1 (1:4000; Sigma; G6046), rabbit anti-G3BP2 (1:4000; Invitrogen, PA5-101815), and rabbit anti-CAPRIN1 (1:4000; Sigma; HPA018126). After washing with TBST, membranes were incubated with HRP-conjugated anti-rabbit IgG secondary antibody (1:5000; Jackson ImmunoResearch; 711-005-152) diluted in 5% BSA for 1 h at RT. Following additional washes, immunoreactive bands were visualized using Clarity™ Western ECL substrate (Bio-Rad; 1705060) using Biorad ChemiDoc^TM^ MP Imaging System.

### RNA immunoprecipitation (RIP) from whole neurons and axonal fractions

RIP was performed using lysates prepared from both whole primary rat cortical neurons and isolated axonal fractions. Axons were isolated using the method described in our previous manuscript (*83*). At 9 *DIV*, neurons/axon fractions were washed twice with PBS and lysed by rotation in RIP buffer containing 150 mM NaCl, 10 mM Tris-HCl, 1 mM EDTA, 1 mM EGTA, 1% Triton X-100, 0.5% NP-40, and 1 mM DTT, supplemented with 1x protease inhibitor cocktail and RNasin Plus (Invitrogen; 10777019). Lysates were clarified by centrifugation at 10,000 x *g* for 15 min at 4 °C. The cleared supernatants were incubated with rabbit anti-G3BP1 antibody (1 µg; Sigma; G6046) for 2 h at 4 °C, followed by incubation with Protein A/G magnetic beads for an additional 2 h with rotation to precipitate RNA-protein complexes. Bead-only samples processed in parallel without primary antibody served as negative controls. Beads were washed eight times with cold RIP buffer, and the bound RNAs were purified.

### RNA isolation and reverse transcriptase digital droplet PCR (RT-ddPCR)

Total RNA from input samples and immunoprecipitates was isolated using TRIzol reagent (Invitrogen; 15596026) and chloroform (VWR; 0757), followed by isopropanol precipitation and washing with 75% ethanol. Purified RNA was resuspended in RNase-free water, and RNA concentration was quantified using the RiboGreen RNA assay (Invitrogen; R11490) according to the manufacturer’s instructions. Reverse transcription was performed using the LunaScript® RT SuperMix Kit (NEB; E3010L). RT-ddPCR was carried out using the QX200™ ddPCR™ EvaGreen Supermix (Bio-Rad; 1864034), and droplets were generated and read using the QX200™ droplet generator and droplet reader, respectively (Bio-Rad). Custom transcript-specific primer sets were used for RT-ddPCR amplification as listed below.

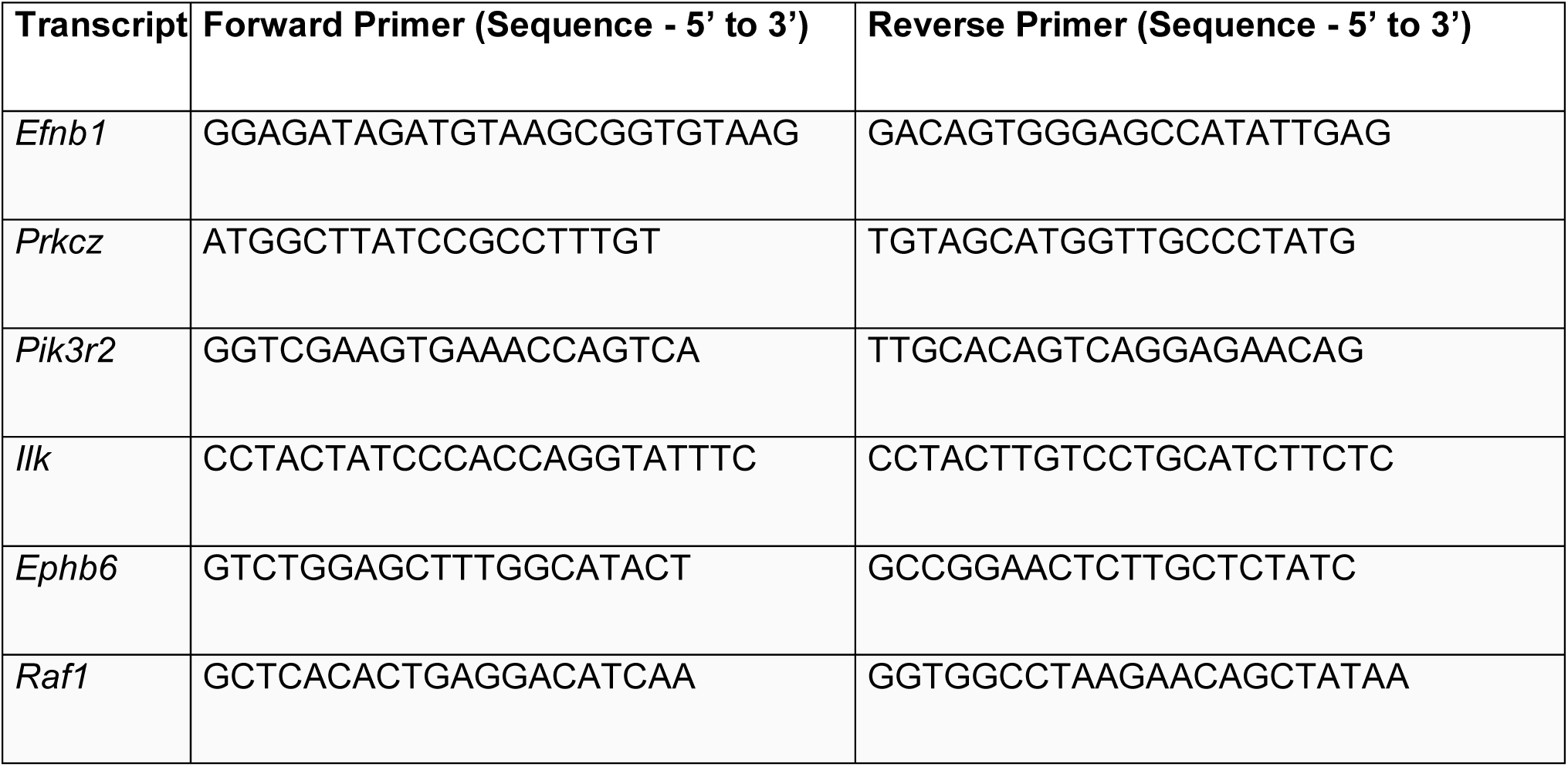
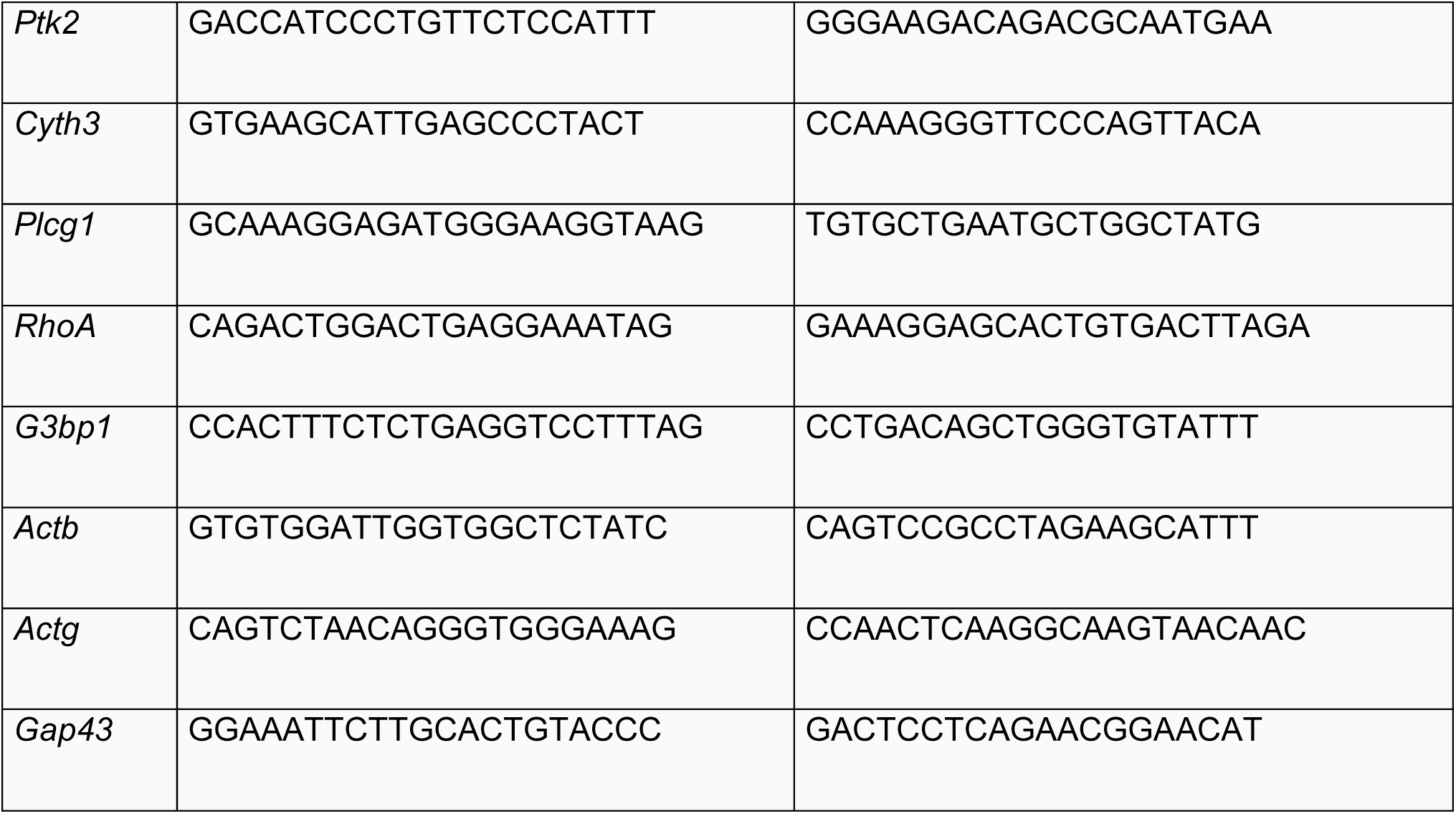

### RNA-Seq Analysis

RNA-seq data from G3BP1 and control (non-specific IgG) RIP and corresponding input samples prepared from the axonal compartments of embryonic rat cortical neurons were analyzed using a standard analysis pipeline (https://github.com/icnn/RNAseq-PIPELINE.git). Briefly, reads were aligned to the rat reference genome (Rnor_6.0/rn6) using STAR (v2.5.3) with default parameters. Uniquely mapped reads were assigned to rat Ensembl gene models and counted with HTSeq (v0.11.3). Genes with low expression were filtered out (retaining genes with counts-per-million ≥ 0.1 in 6 samples), and the retained counts were normalized by the trimmed mean of M-values (TMM) method. Differential enrichment was assessed in edgeR (v3.30.0) by fitting a negative-binomial generalized linear model and evaluating contrasts (notably G3BP1 IP vs. control IP for each genotype) with likelihood-ratio tests; to account for replicate-associated batch variation identified by principal component analysis, replicate batch was included as a covariate in the model. P-values were adjusted for multiple testing using the Benjamini–Hochberg false discovery rate (FDR), and transcripts with FDR < 0.1 and a positive log2 fold change were considered significantly enriched in the G3BP1 immunoprecipitate. Gene Ontology enrichment of differentially associated transcripts was performed using DAVID, and rank–rank hypergeometric overlap (RRHO) analysis was used to compare the concordance of enrichment signatures across conditions.

### Puromycinylation

The puromycinylation assay was performed as described previously(*27*). Briefly, at 11 *DIV*, cells were starved using Neurobasal medium with 1x L-Glutamine for 3 h. Three hours after starvation, cells were stimulated with Neurobasal medium supplemented with 1x L-Glutamine, 1x B27, 1 μg/mL laminin, and 100 ng/μL brain-derived neurotrophic factor (BDNF) for 15 min. Anisomycin was added for a total duration of 45 min, starting at 2 h 30 min of starvation, and puromycin was added at the time of stimulation for a total duration of 15 min. Post-stimulation, cells were fixed in 4% paraformaldehyde (PFA) in PBS, and immunofluorescence was performed.

### Immunofluorescence (IF)

For SG assembly and disassembly in COS-7 cells, after fixation, cells were permeabilized with 0.3% Triton X-100 in PBS for 5 min. Primary and secondary antibodies were prepared in 0.1% Triton X-100 with 10% normal donkey serum (NDS). Cells were incubated with the primary antibodies at 4 °C overnight. Primary antibodies used were rabbit anti-HA (1:500, Sigma Aldrich; H6908)/rat anti-HA (1:500, Roche; C755C30), rabbit anti-G3BP1 (1:1000, Sigma Aldrich; G6046). Following primary antibody, cells were washed three times with PBS for 5 min each. Cells were incubated with the secondary antibodies at RT for 1 h. Secondary antibodies used were Cy3 - conjugated donkey anti-rat (1:200; Jackson ImmunoResearch; 712-165-150), FITC-conjugated donkey anti-rabbit (1:200; Jackson ImmunoResearch; 711-095-152) (for cells transfected with CK2α variants), and Cy5-conjugated donkey anti-rabbit (for cells transfected with GFP) (1:200; Jackson ImmunoResearch; 711-175-152). Cells were mounted using mounting media ProLong Diamond with DAPI (Invitrogen; P36962). Visualization was performed using a Zeiss Axio Observer 7 microscope with an Axiocam 820 mono camera.

For E18 primary rat cortical neurons, IF was performed on cells fixed with 4% PFA for 15 min at RT. Following fixation, cells were washed three times with PBS for 5 min each. Cells were permeabilized with 0.3% Triton X-100 for 10 min. Primary and secondary antibodies were prepared using 0.1% Triton X-100 with 5% normal donkey serum (NDS) in PBS. Cells were incubated with primary antibodies at RT for 1 h followed by secondary antibodies at RT for 30 min. Cells were mounted using mounting media ProLong Gold (Invitrogen; P36934) or ProLong Diamond with DAPI. Visualization was performed using a Zeiss Axio Observer 7 microscope with an Axiocam 820 mono camera.

To study phosphorylation of G3BP1 at S149 (G3BP1^PS149^), primary antibodies used were rabbit anti-G3BP1^PS149^ (1:300, Sigma; G8046) mouse anti-SMI312 (1:500, BioLegend; 837904), mouse anti-Tau (1:500, Cell Signaling Technologies; 4019), and chicken anti-MAP2 (1:3000, Abcam; ab5392). Secondary antibodies were as follows: Cy3-conjugated donkey anti-mouse (1:200, Jackson ImmunoResearch; 715-165-150), Cy5-conjugated donkey anti-rabbit (1:200, Jackson ImmunoResearch; 711-175-152), and Alexa Fluor 405-conjugated goat anti-chicken (1:200, Invitrogen; A48260). For detection of GFP/HA-tagged constructs, pre-labeling was performed using the Zenon IgG labeling kit (Invitrogen; Z25302) for rabbit anti-GFP (1:500; Sigma; SAB4301138) for cells transfected with GFP and rabbit anti-HA (1:500, Sigma Aldrich; H6908) for cells transfected with plasmids encoding CK2α variants.

For polarity and granularity in rat cortical neurons, primary antibodies used were chicken anti-MAP2 (1:3000, Abcam; ab5392), mouse anti-SMI312 (1:500, BioLegend; 837904), mouse anti-Tau (1:500, Cell Signaling Technologies; 4019), rabbit anti-GFP (1:500, Sigma; SAB4301138), rabbit anti-G3BP1 (1:200, Sigma; G6046), and rabbit anti-G3BP2 (1:200, Invitrogen, PA5-101815). Secondary antibodies were as follows: Cy3-conjugated donkey anti-mouse (1:200, Jackson ImmunoResearch; 715-165-150), Cy5-conjugated donkey anti-rabbit (1:200, Jackson ImmunoResearch; 711-175-152), and Alexa Fluor 405-conjugated goat anti-chicken (1:200, Invitrogen; A48260). For detection of GFP/HA-tagged constructs, pre-labeling was performed using the Zenon IgG labeling kit for rabbit anti-GFP (1:500, Sigma; SAB4301138) for cells transfected with GFP and rabbit anti-HA (1:500, Sigma Aldrich; H6908) for cells transfected with plasmids encoding CK2α variants.

For puromycinylation in rat cortical neurons, primary antibodies used were chicken anti-MAP2 (1:3000, Abcam; ab5392), mouse anti-SMI312 (1:500, BioLegend; 837904), mouse anti-Tau (1:500, Cell Signaling Technologies; 4019), rabbit anti-GFP (1:500, Sigma; SAB4301138), or rabbit anti-HA (1:500, Sigma Aldrich; H6908). Secondary antibodies were as follows: FITC-conjugated donkey anti-mouse (1:200, Jackson ImmunoResearch; 715-095-150), Cy3-conjugated donkey anti-rabbit (1:200, Jackson ImmunoResearch; 711-165-152), and Alexa Fluor 405-conjugated goat anti-chicken (1:200, Invitrogen; A48260).

To label dendrites and synapses, primary antibodies used were chicken anti-MAP2 (1:3000, Abcam; ab5392), mouse anti-PSD95 (1:50, DSHB; K28/43-s), and rabbit anti-synaptophysin (1:100, Abcam; ab14692). Secondary antibodies were as follows: Cy3-conjugated donkey anti-mouse (1:200, Jackson ImmunoResearch; 715-165-150), Cy5-conjugated donkey anti-rabbit (1:200, Jackson ImmunoResearch; 711-175-152), and Alexa Fluor 405-conjugated goat anti-chicken (1:200, Invitrogen; A48260). For detection of GFP/HA-tagged constructs, pre-labeling was performed using the Zenon IgG labeling kit (Invitrogen; Z25302) for rabbit anti-GFP (1:500; Sigma; SAB4301138) for cells transfected with GFP and rabbit anti-HA (1:500; Sigma Aldrich; H6908) for cells transfected with plasmids encoding CK2α variants.

For the detection of G3BP1 granularity and G3BP1^PS149^ *in vivo*, free-floating sections were immunostained as described previously (*84*). Briefly, sections were incubated in PBS for 10 min at RT, followed by three washes with freshly made 20 mM glycine for 10 min each at RT. Sections were then incubated with 0.25 M NaBH_4_ for 30 min at RT, changing every 10 min. The sections were then rinsed with wash buffer containing 20 mM glycine and 0.1% Tween-20 in PBS. Next, sections were permeabilized in 0.2% Triton X-100 made in PBS for 15 min at RT. Sections were then blocked in PBS containing 20 mM glycine, 0.1 % Tween-20, and 10% NDS for 1 h at RT. Sections were incubated with the primary antibodies in blocking buffer at 4 °C overnight. The next day, sections were washed three times for 5 min each at RT with wash buffer containing 20 mM glycine and 0.1% Tween-20 in PBS. The sections were incubated with the secondary antibodies in blocking buffer for 1 h at RT. Primary antibodies used were chicken anti-MAP2 (1:2000, Abcam; ab5392), mouse anti-SMI312 (1:500; BioLegend; 837904), mouse anti-Tau (1:500, Cell Signaling Technologies; 4019), rabbit anti-G3BP1 (1:100, Sigma Aldrich; G6046), and rabbit anti-G3BP1^PS149^ (1:300; Sigma Aldrich; G8046). Secondary antibodies were as follows: FITC-conjugated donkey anti-mouse (1:200, Jackson ImmunoResearch; 715-095-150), Cy3-conjugated donkey anti-rabbit (1:200, Jackson ImmunoResearch; 711-165-152), and Cy5-conjugated donkey anti-chicken (1:200, Jackson ImmunoResearch; 703-175-155). The sections were washed thrice for 5 min each at RT with wash buffer containing 20 mM glycine and 0.1% Tween-20 in PBS. Sections were mounted on subbed slides using mounting media ProLong Diamond with DAPI (Invitrogen). Visualization was performed using a Zeiss LSM980 Confocal Microscope with Airyscan 2.

For hiPSC-derived iNeurons, cells plated on coverslips in a 12-well plate were grown to 7 *DIV* and then fixed with 4% PFA and immunostained, similar to the primary rat cortical neurons. Primary antibodies used were chicken anti-MAP2 (1:200, Aves; MAP), rabbit anti-G3BP1 (1:100, Sigma Aldrich; G6046), and Hoechst 33258 (1 µg/mL, Invitrogen; H3569). Secondary antibodies used were Alexa Fluor 647 anti-rabbit (1:500, ThermoFisher; A21245) and Alexa Fluor 488 anti-chicken (1:500, ThermoFisher; A11039).

### Microelectrode arrays (MEA)

Sterile MEAs (HBio Multichannel Systems, 60MEA200/10iR-Ti) consisting of 59 working electrodes and 1 reference electrode were treated with oxygen plasma, followed by an overnight treatment with 100 µg/mL poly-D-lysine and a 2 h treatment with 10 µg/mL laminin. Cortical cells (5 × 10^6^) were nucleofected with 1 μg each of CK2α^WT^, CK2α^K198R^ or GFP plasmids and co-transfected with 1 μg mCherry plasmid as described above and plated at a density of approximately 2,100 cells/mm^2^ on MEAs.

Extracellular recordings were obtained at 7, 14, and 21 *DIV* as previously described(*85, 86*). Conditioned medium was replaced with recording buffer containing 144 mM NaCl, 10 mM KCl, 1 mM MgCl_2_, 2 mM CaCl_2_, 10 mM HEPES, 2 mM Na-pyruvate, 10 mM glucose, and pH = 7.4, and cultures were stabilized for 5 min at 37 °C and 5% CO_2_. Recordings were collected using the MEA2100-Lite rig and Multi Channel Experimenter Software. The MEA2100-Lite rig head stage was maintained at 37 °C using a TC02 temperature control unit and TCX-Control software. Five minutes of data were collected with a sample rate of 20 kHz. Raw data were exported for analysis in MATLAB.

Raw data were filtered with a fourth-order Butterworth bandpass filter (20-2000 Hz) and a comb filter to remove 60 Hz noise and its harmonics(*85*). The signal was divided into 10 s blocks for spike detection. Spikes were calculated via amplitude thresholding (*87*), where a spike was defined as an event where the filtered signal amplitude exceeded five times the estimated standard deviation. Each electrode and each 10 s block had its own signal threshold. The minimum interspike interval (ISI) was designated for 2 ms to ensure that the same spike was not counted twice. Spike rate for individual electrodes and the overall network were defined as spikes per minute of recording. Fano factor, a measure of variability in a neuron’s firing pattern, was calculated by binning spikes into 100 ms bins and quantifying the ratio of the variance of spiking between bins to the mean of spike counts between bins:

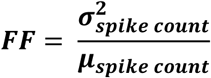

### Image acquisition and analyses

All images were captured and analyzed by epifluorescent microscopy (Zeiss Axio Observer 7 motorized inverted microscope with Axiocam 820 mono camera) or confocal microscopy (Zeiss LSM980 Confocal Microscope with Airyscan 2). For quantitation between samples, imaging parameters were matched for exposure, gain, offset, and post-processing(*27*). All analyses were performed in a blinded manner. All experiments were repeated with at least three independent biological replicates, unless specified in the figure legends.

To calculate the percentage of COS-7 cells with SGs, the number and size of SGs were quantified in ImageJ using the ComDet v.0.5.6 plugin, with consistent detection thresholds across all samples. A size cutoff of 0.4 µm² was applied for SG detection. For each cell, the number and size of G3BP1 granules were measured, and the percentage of transfected cells containing detectable G3BP1 granules was manually calculated.

For calculating axonal and dendritic G3BP1 and G3BP2 granules, 15-25 µm axonal segments ≥ 100 µm away from the soma and 10-15 µm dendritic segments ≥ 50 µm away from the soma were considered. The ImageJ segmented line tool was used to track the axons (SMI312/Tau^+^) and dendrites (MAP2^+^), and SMI312/Tau or MAP2 overlapping SG protein signals were projected as a separate channel. ImageJ particle analyzer was used to quantify the number and size of the granules. Only granules bigger than 0.01 µm^2^ were considered for analysis.

For puromycinylation and G3BP1^PS149^ quantifications, soma and 15-25 µm axonal segments ≥ 100 µm away from the soma, and 10-15 µm dendritic segments ≥ 50 µm away from the soma were considered(*27*). The ImageJ polygon tool was used to mark the soma, and the segmented line tool was used to mark the axons and dendrites, tracking the SMI312/Tau and MAP2 signal, respectively. Mean fluorescence intensity was quantified using ImageJ for puromycinylation and G3BP1^PS149^ signals.

For neurite outgrowth in primary rat cortical neurons, images from 11 *DIV* cortical cultures were analyzed for neurite length and branching complexity using WIS-Neuromath(*88*) with SMI312/Tau and MAP2 immunostained cultures(*27*).

For calculation of the number of synapses in primary rat cortical neurons, images from 21 *DIV* cortical cultures were analyzed by assessing the colocalization of presynaptic (synaptophysin) and postsynaptic (PSD95) markers using ImageJ with ComDet v.0.5.6 plugin. Specifically, puncta corresponding to PSD95 and synaptophysin were detected and quantified independently in their respective channels using identical detection parameters across conditions. Colocalization was defined as puncta from the PSD95 and synaptophysin channels whose centroids were within a predefined maximum distance (≤4 pixels). The number of colocalized puncta was quantified per image and normalized to dendritic length, as indicated.

For calculating axonal and dendritic G3BP1 and G3BP1^PS149^ intensity in mouse brain sections, the ImageJ colocalization plugin was first used to extract G3BP1 and G3BP1^PS149^ signals, respectively, that overlap with the axonal marker SMI312/Tau or dendritic marker MAP2 in the brain sections to extract the ‘axon-only’ or ‘dendrite-only’ signal projected as a separate channel(*27*). The colocalized signal was normalized to the SMI312/Tau signal for axonal G3BP1 and G3BP1^PS149^ intensity and to MAP2 for dendritic G3BP1 and G3BP1^PS149^ intensity.

For calculating the number and size of axonal and dendritic G3BP1 granules in mouse brain sections, the ImageJ segmented line tool was first used to track the axons (SMI312/Tau^+^) and dendrites (MAP2^+^), followed by the ImageJ ComDet v.0.5.6 plugin with consistent detection thresholds (20 for G3BP1 and 100 for SMI/Tau and MAP2) across all samples.

To examine growth rate in hiPSC-derived iNeurons, cells were plated in a 96-well plate and allowed to attach to the bottom. Cells were imaged daily using an Incucyte S3 (Sartorius) and its Neurotrack package, with cell and neurite masks.

For neurite morphology analysis in hiPSC-derived iNeurons, images from 7 *DIV* cultures were analyzed for neurite length and branching complexity using WIS-Neuromath(*88*) in MAP2-immunostained cultures(*27*).

To calculate G3BP1 and G3BP1^PS149^ intensities in hiPSC-derived iNeurons, the ImageJ Max Intensity Projection tool was first used to extract the G3BP1 signal from whole neurons. The G3BP1 signal was then normalized to the MAP2 signal for G3BP1 intensity.

For calculating axonal G3BP1 granules in the hiPSC-derived iNeurons, 15-25 µm neurite segments ≥ 50 µm away from the soma were considered. The ImageJ segmented line tool was used to track the primary neurites using MAP2, and MAP2 overlapping G3BP1 signal was projected as a separate channel. Image J particle analyzer was used to quantify the number and size of the granules. Only granules bigger than 0.01 µm^2^ were considered for analyses.

## QUANTIFICATION AND STATISTICAL ANALYSIS

Prism (GraphPad) software package was used for statistical analyses. Normality tests were performed on all data to determine whether they were normally distributed. One-way ANOVA, two-way ANOVA, or an equivalent nonparametric test was used to compare means of >2 independent groups, and Student’s *t*-test or Mann-Whitney test was used to compare between 2 groups with appropriate post-hoc tests as stated in the figure legends. *P* values of ≤0.05 were considered statistically significant.

## RESOURCE AVAILABILITY

### Lead contact

Requests for further information and resources should be directed to and will be fulfilled by the lead contact, Dr. Pabitra K. Sahoo.

### Materials availability

The data that support the findings of this study are all provided with this manuscript. Reagents generated in this study will be available through materials transfer agreements (MTAs) with Rutgers University-Newark.

## ACKNOWLEDGMENTS

This work was supported by grants from the Merkin Peripheral Neuropathy and Nerve Regeneration Center, Rutgers University Newark startup fund (to P.K.S.), the New Jersey Autism Center of Excellence (to M.D.), the New Jersey Commission on Brain Injury Research (Fellowship #CBIR24FEL009 to B.J.V. and Grant #CBIR24IRG005 to B.L.F., #CBIR26IRG029 to P.K.S.), and NIH/NINDS (Grant #1R01NS135406-01A1 to B.L.F.), CSNK2A1 foundation research grant (to H.R.), Natural Science and Engineering Research Council of Canada (NSERC)-Postdoctoral Fellowship (to K.S.), The Eagle’s Autism Foundation (to J.P.), and Dr. Miriam and Sheldon G. Adelson Medical Research Foundation (to D.H.G.). We thank Prof. Jeffery L. Twiss for the critical reading of the manuscript.

## AUTHOR CONTRIBUTIONS

Conceptualization – M.A., M.D., S.G., P.K.S.

Methodology – M.A., M.D., S.G., Y.B., B.J.V., K.S., K.R., N.H.H., P.N., N.S., A.L., R.K., P.K.S.

Investigation – M.A., M.D., S.G., Y.B., B.J.V., K.S., K.R., N.H.H., P.N., N.S., A.L., R.K., P.K.S.

Visualization – M.A., M.D., S.G., Y.B., B.J.V., K.S., P.K.S.

Funding acquisition – B.L.F., J.P., D.H.G., H.R., P.K.S.

Project administration – P.K.S.

Supervision – B.L.F., J.P., D.H.G., H.R., P.K.S.

Writing, original draft – M.A., M.D., S.G., Y.B., B.J.V., K.S., R.K., B.L.F., P.K.S.

Writing, review & editing – M.A., M.D., S.G., Y.B., B.J.V., K.S., B.L.F., H.R., P.K.S.

## SUPPLEMENTAL INFORMATION

### Document S1

**Figures S1–S7, Movie S1, and Table 1 (see below)**

**Figure S1:**
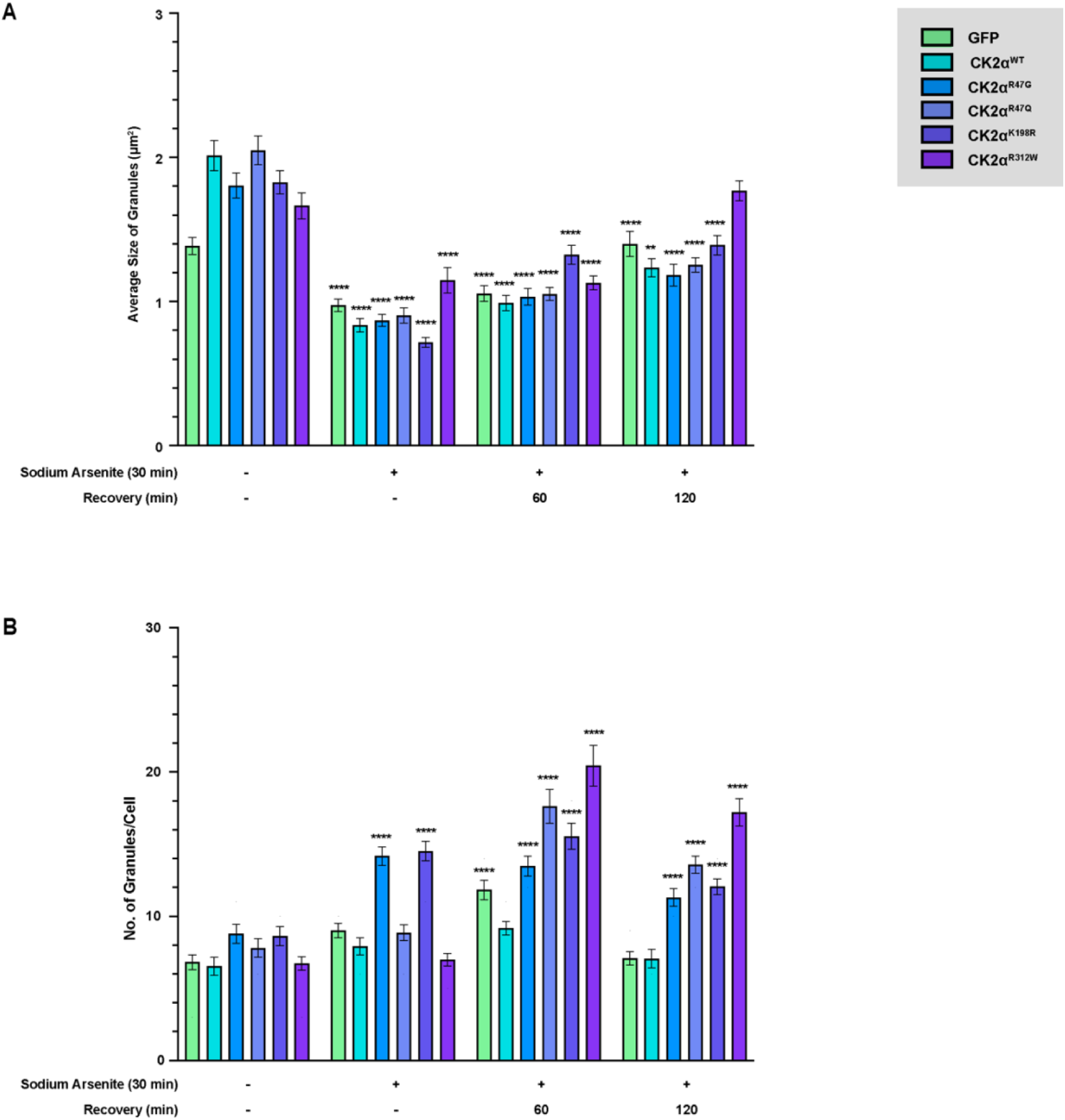
COS-7 cells expressing CK2α mutants fail to disassemble the stress granules. ***A-B,*** Quantifications of the average granule size **(A)** and the number of G3BP1 granules/cell **(B)** in COS-7 cells transfected with either GFP, CK2α^WT^, or CK2α mutants are shown as mean ± SEM. n ≥ 65 cells in **(A)** and n ≥ 434 granules in **(B)** across three biological replicates; *p ≤ 0.05, **p ≤ 0.01, ***p ≤ 0.001, ****p ≤ 0.0001 by Kruskal-Wallis test with Dunn’s multiple comparisons vs. no stress GFP control.

**Figure S2:**
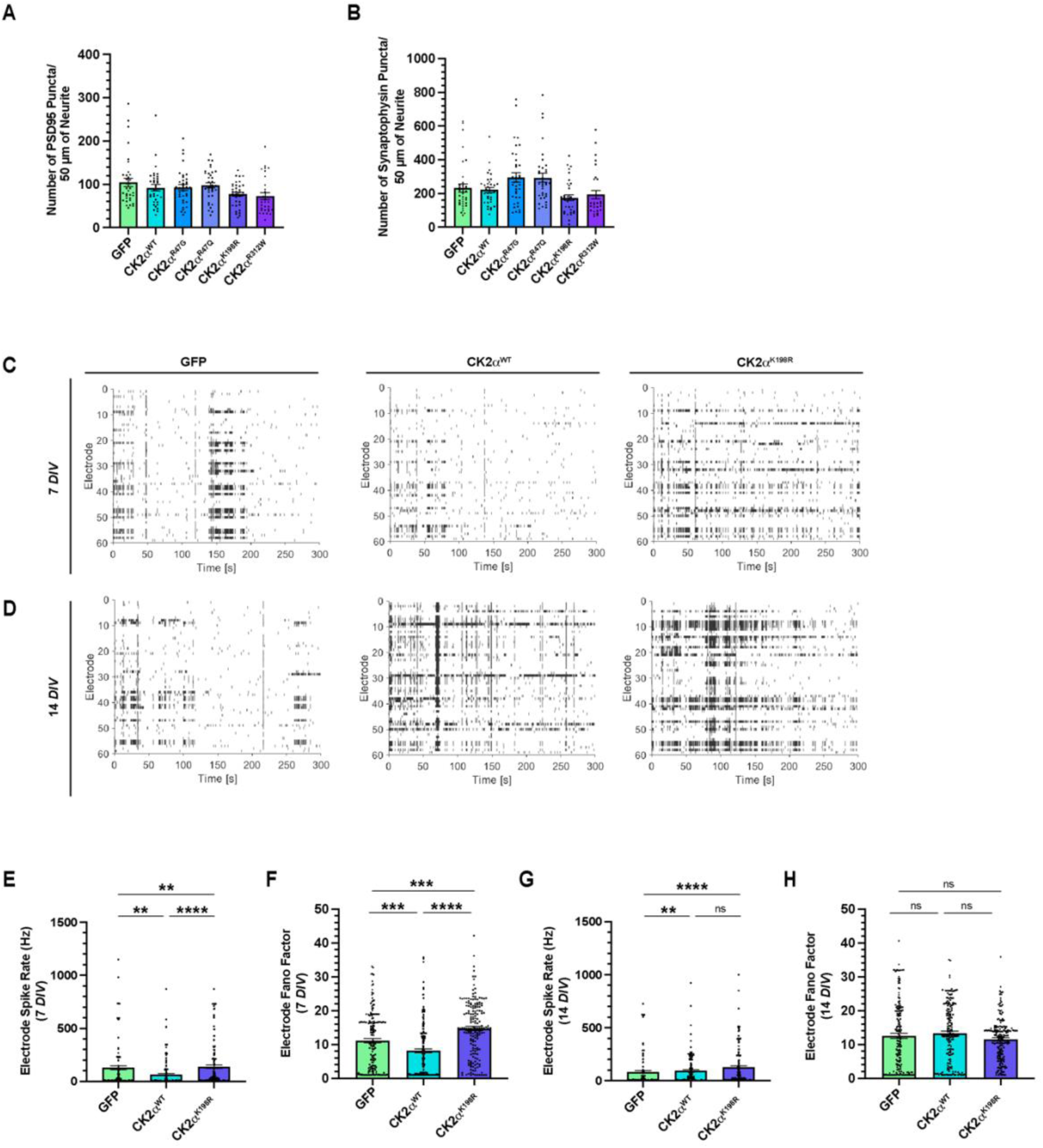
OCDNS-associated CK2α^K198R^ mutant leads to hyperactivity of developing rat cortical neurons during maturation. ***A-B,*** Quantification of the number of PSD95 puncta **(A)** and the number of synaptophysin puncta **(B)** is shown as mean ± SEM. n ≥ 30 neurites across three biological replicates; ns by Kruskal-Wallis test with Dunn’s multiple comparisons. ***C-D,*** Raster plots for activity detected using MEAs for embryonic rat cortical neurons expressing either GFP, CK2α^WT^, or CK2α^K198R^ cultured for 7 **(C)** and 14 **(D)** *DIV* are shown. ***E-H,*** Quantifications of electrode spike rate **(E)** and electrode Fano factor **(F)** for embryonic rat cortical neurons expressing either GFP, CK2α^WT^ or CK2α^K198R^ cultured for 7 *DIV* and electrode spike rate **(G)** and electrode Fano factor **(H)** for those cultured for 14 *DIV* are shown as mean ± SEM. n = recording from 177 electrode across three technical replicates; **p ≤ 0.01, ***p ≤ 0.001, ****p ≤ 0.0001 by Kruskal-Wallis test with Dunn’s multiple comparison.

**Figure S3:**
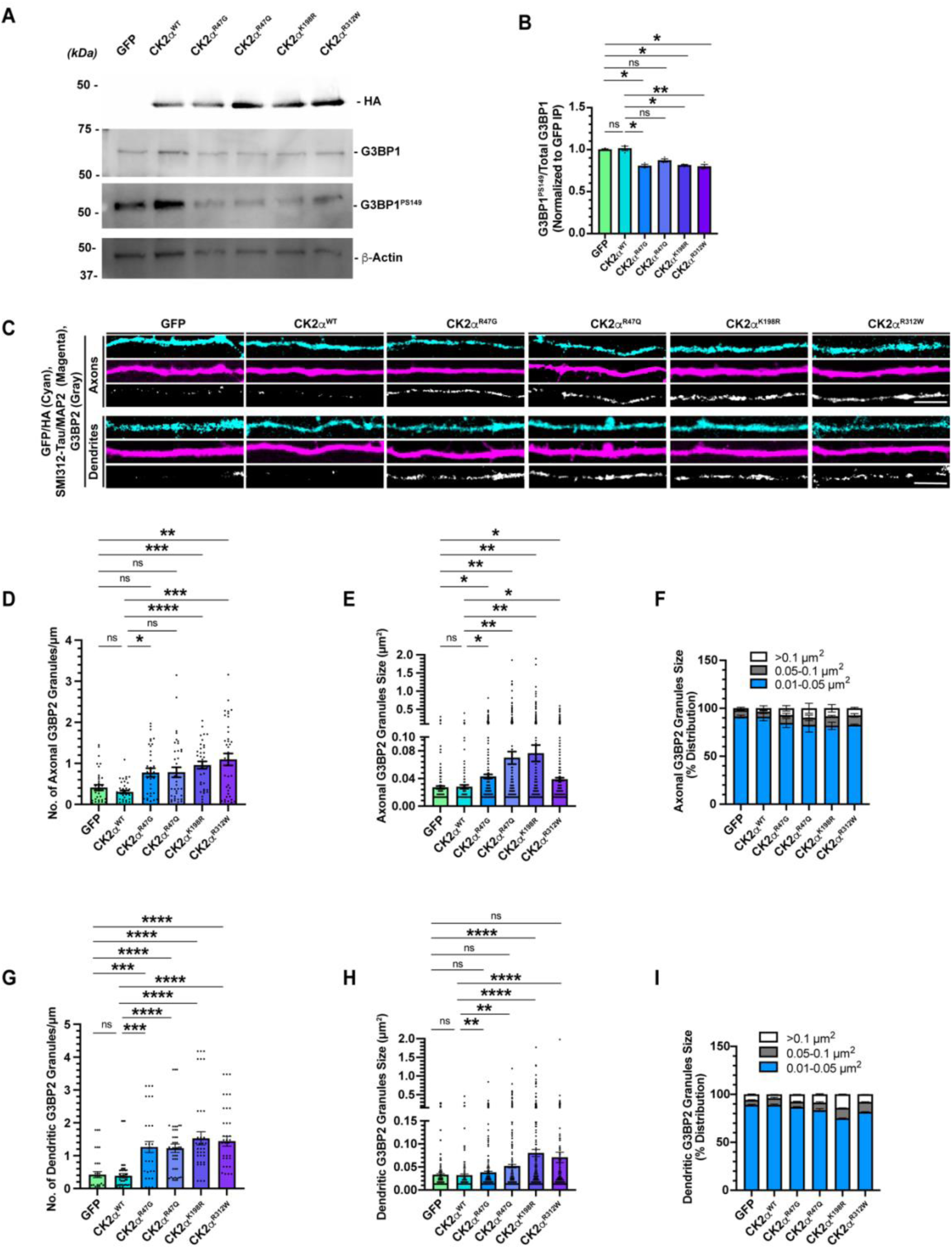
OCNDS-associated CK2α mutants increase the number and size of axonal and dendritic G3BP2 granules in developing rat cortical neurons. ***A-B,*** Immunoblot shows the levels of pan-G3BP1 and G3BP1^PS149^ in E18 rat cortical neurons, cultured for 9 *DIV,* expressing either GFP, CK2α^WT^, or CK2α mutants. β-Actin was used as a loading control (**A**). Densitometric quantifications of G3BP1^PS149^ level normalized to pan-G3BP1 are shown as mean ± SEM across three biological replicates (**B**); *p ≤ 0.05, **p ≤ 0.01 by Friedman test with Dunn’s multiple comparisons. ***C,*** Representative confocal images of axons (top) and dendrites (bottom) of E18 rat cortical neurons expressing either GFP, CK2α^WT^, or CK2α mutants cultured for 11 *DIV* and immunostained for SMI312/Tau, G3BP2, and GFP or HA are shown (Scale bar, 5 µm). ***D-F,*** Quantifications of the number of axonal G3BP2 granules/μm **(D)**; n ≥ 36 axons, average granule size **(E)**; n ≥ 181 granules, and percentage distribution of granules as per size **(F)** are shown as mean ± SEM across three biological replicates; *p ≤ 0.05, **p ≤ 0.01, ***p ≤ 0.001, ****p ≤ 0.0001 by Kruskal-Wallis test with Dunn’s multiple comparisons. ***G-I,*** Quantifications of the number of dendritic G3BP2 granules/μm **(G)**; n ≥ 33 dendrites, average granule size **(H)**; n ≥ 318 granules, and percentage distribution of granules as per size **(I)** are shown as mean ± SEM across three biological replicates; **p ≤ 0.01, ***p ≤ 0.001, ****p ≤ 0.0001 by Kruskal-Wallis test with Dunn’s multiple comparisons.

**Figure S4:**
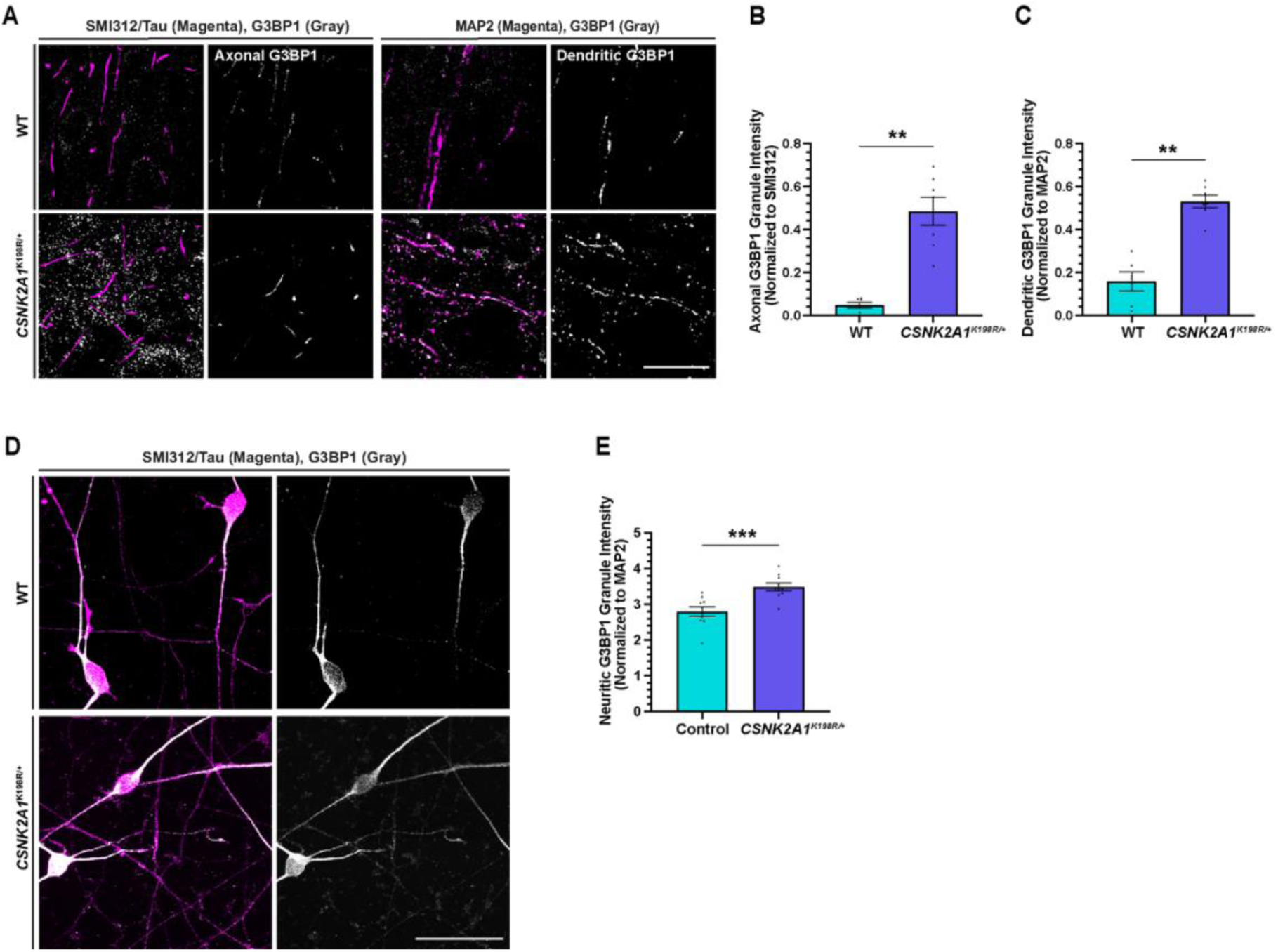
G3BP1 granule dynamics are impaired in mouse and human OCNDS models. ***A,*** Representative confocal images of axons (left) and dendrites (right) of cortices from coronal sections of adult *CSNK2A1*^K198R/+^ or WT mouse brain immunostained for SMI312/Tau, MAP2, and G3BP1 are shown (Scale bar, 50 µm)**. *B-C,*** Quantifications of G3BP1 granule intensity levels colocalizing with and normalized to SMI312/Tau for axons **(B)** and MAP2 for dendrites **(C)** are shown as mean ± SEM. n ≥ 6 images across five animals from each genotype; **p ≤ 0.01 by Mann Whitney. ***D,*** Representative confocal images with maximum intensity projection of *CSNK2A1*^K198R/+^ iNeurons immunostained for MAP2 and G3BP1 are shown (Scale bar, 5 µm)**. *E,*** Quantification of total G3BP1 granule intensity levels colocalizing with and normalized to MAP2 is shown as mean ± SEM. n ≥ 10 images across three biological replicates; ***p ≤ 0.001 by Mann Whitney.

**Figure S5:**
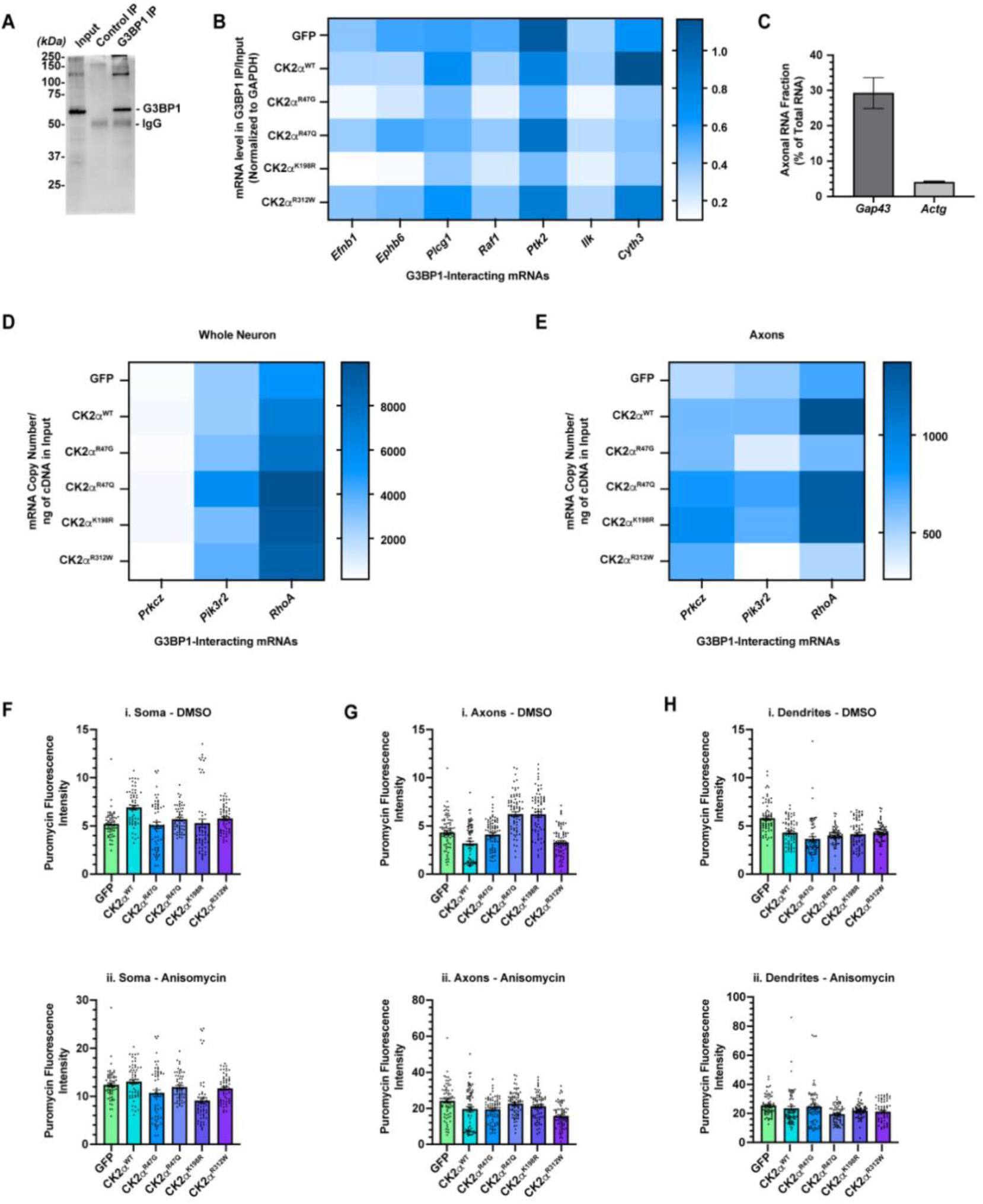
Expression of specific OCNDS-associated CK2α mutants reduce axonal mRNA levels. ***A,*** Immunoblot showing immunoprecipitation of G3BP1 from E18 rat cortical neurons cultured for 9 *DIV*. ***B,*** Heat map shows the quantifications of mRNA levels in the indicated expression conditions from whole neurons following G3BP1 immunoprecipitation and RT-ddPCR analysis are shown as mean across three biological replicates. ***C,*** A higher percentage of *Gap43* mRNA vs. *Actg* mRNA present in axons relative to their whole neuron level is shown as mean ± SEM across three biological replicates, showing enrichment of axonal preparations. ***D-E,*** Heat maps show the mRNA copy numbers in the inputs of whole neurons (**D**) and axons (**E**) are shown as the mean across three biological replicates. ***F,*** Quantifications of puromycin levels in the soma when treated with DMSO **(Fi)**, and when treated with puromycin and anisomycin **(Fii)** are shown as mean ± SEM. n ≥ 60 soma across three biological replicates. ***G,*** Quantifications of puromycin levels in the axons when treated with DMSO **(Gi)**, and when treated with puromycin and anisomycin **(Gii)** are shown as mean ± SEM. n ≥ 72 axons across three biological replicates. ***H,*** Quantification of puromycin levels in the dendrites when treated with DMSO **(Hi)**, and when treated with puromycin and anisomycin **(Hii),** is shown as mean ± SEM. N ≥ 66 dendrites across three biological replicates.

**Figure S6:**
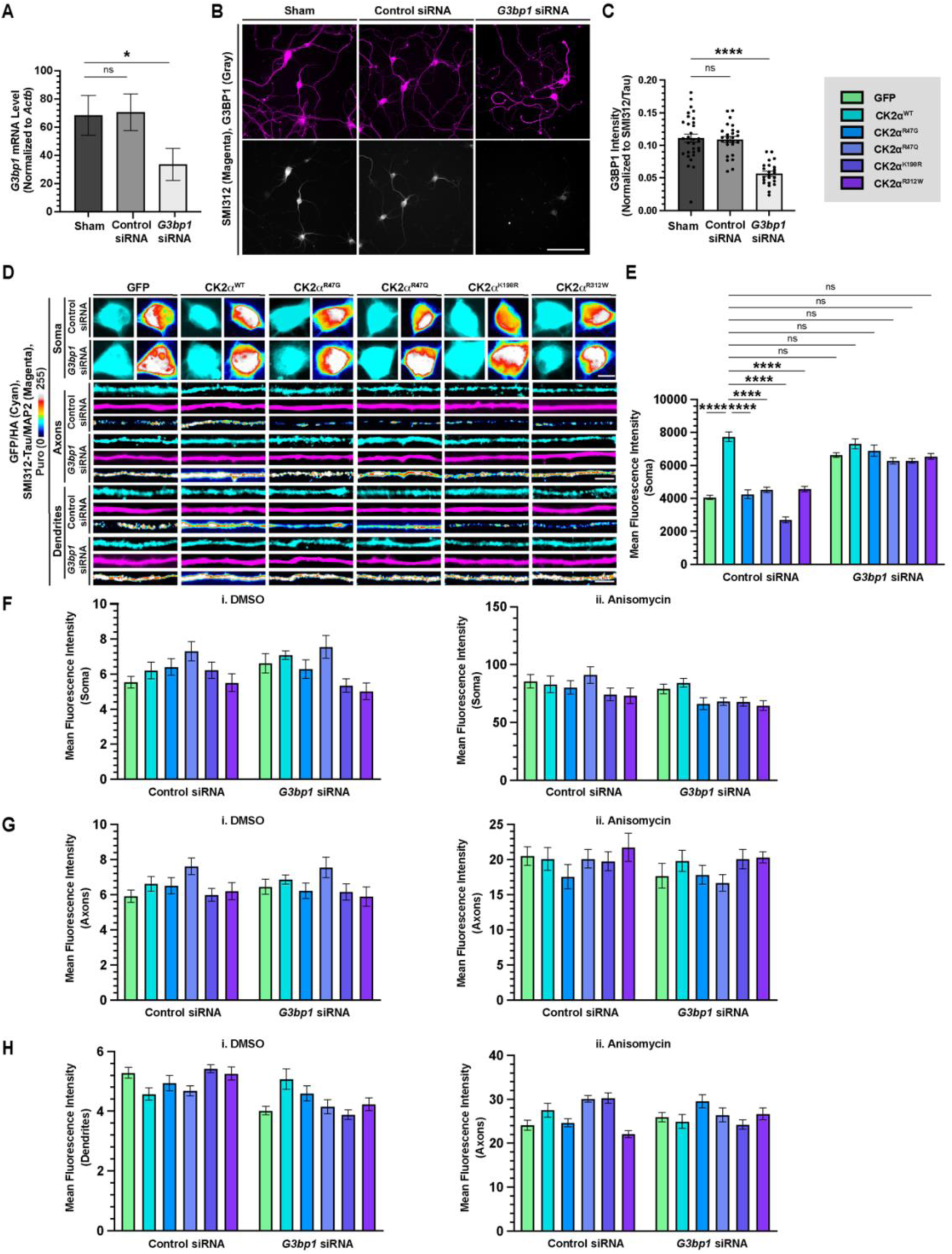
Knocking down G3bp1 rescues defects in protein synthesis. ***A,*** Quantification of the number of *G3bp1* mRNA copies detected using RT-ddPCR in E18 rat cortical neurons cultured for 9 *DIV* and treated with control or *G3bp1* siRNA for final 3 *DIV* is shown as mean ± SEM across three biological replicates; *p ≤ 0.05 by Kruskal-Wallis with Dunn’s multiple comparisons. ***B,*** Representative epifluorescent images of E18 rat cortical neurons transfected with GFP, cultured for 11 *DIV,* treated with *control or G3bp1* siRNA on 7 *DIV* for 4 *DIV,* and immunostained for SMI312/Tau and G3BP1 *are* shown (Scale bar, 100 µm). ***C,*** Quantification of total G3BP1 intensity level colocalizing with and normalized to SMI312/Tau is shown as mean ± SEM. n ≥ 23 images across three biological replicates; ns, ****p ≤ 0.0001 by ordinary one-way ANOVA with Dunnett’s multiple comparisons. ***D,*** Representative epifluorescent images of soma (top), axons (middle), and dendrites (bottom) of E18 rat cortical neurons expressing either GFP, CK2α^WT^ or CK2α mutants cultured for 11 *DIV*, treated with *control or G3bp1* siRNA on 7 *DIV* for 4 *DIV,* treated with puromycin, and immunostained for puromycin, SMI312/Tau or MAP2 and GFP or HA are shown (Scale bar, 10 µm for soma and 5 µm for axons and dendrites). ***E,*** Quantifications of puromycin levels in the soma when treated with puromycin is shown as mean ± SEM. n ≥ 43 soma across three biological replicates. ***F,*** Quantifications of puromycin levels in the soma when treated with DMSO **(Fi)** and anisomycin **(Fii)** are shown as mean ± SEM. n ≥ 45 soma across three biological replicates. ***G,*** Quantifications of puromycin levels in the axons when treated with DMSO **(Gi)**, and puromycin plus anisomycin **(Gii)** are shown as mean ± SEM. n ≥ 59 axons across three biological replicates. ***H,*** Quantifications of puromycin levels in the dendrites when treated with DMSO **(Hi)**, and puromycin plus anisomycin **(Hii)**, are shown as mean ± SEM. n ≥ 58 dendrites across three biological replicates.

**Figure S7:**
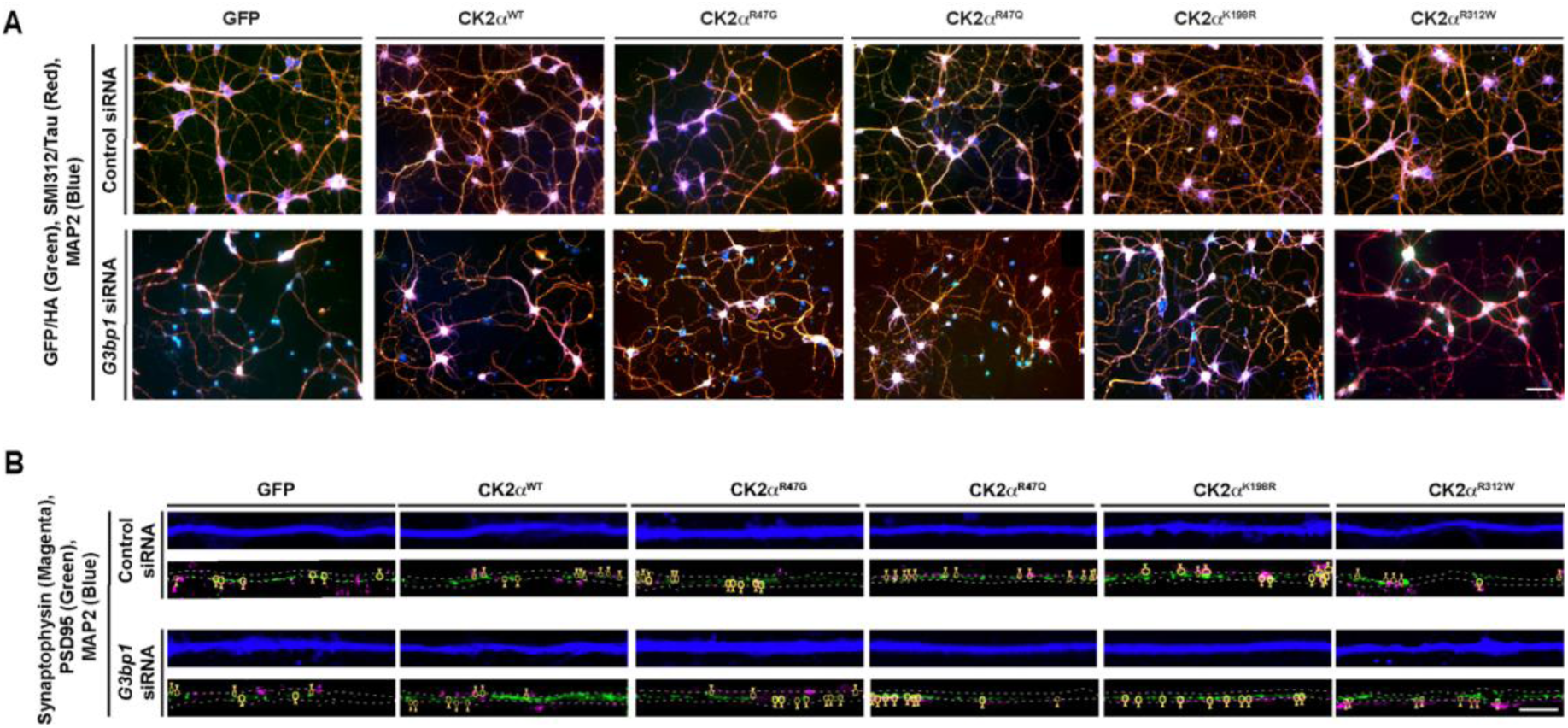
Knocking down G3bp1 partially rescues defects in neurite growth and synaptogenesis. ***A,*** Representative epifluorescent images of E18 rat cortical neurons expressing either GFP, CK2α^WT^ or CK2α mutants, cultured for 11 *DIV,* treated with *control or G3bp1* siRNA on 7 *DIV* for 4 *DIV*, and immunostained for SMI312/Tau, MAP2, and GFP or HA are shown (Scale bar, 100 µm). ***B,*** Representative confocal images of embryonic rat cortical neurons expressing either GFP, CK2α^WT^ or CK2α mutants cultured for 21 *DIV*, treated with *control or G3bp1* siRNA on 13 and 17 *DIV* for a total of 8 *DIV*, and immunostained for MAP2, Synaptophysin, and PSD95 are shown (Scale bar, 5 µm).

**Video S1. FRAP Movies, related to Figure 1C-D**

Videos showing GFP-G3BP1 granules in neurons co-expressing mCherry control, mCherry-CK2α^WT^, or mCherry-CK2α^K198R^. Red dashed circles indicate a photobleached GFP-G3BP1 granule. ROIs monitored over time; refer to Figure 1 for quantitation. Time stamps are shown in seconds; scale bar = 5 μm.

**Table 1. Axonal G3BP1 RIP-seq Dataset, related to Figure 5**

Differential enrichment analysis of RNAs recovered by axonal G3BP1 RNA immunoprecipitation sequencing compared with control immunoprecipitation. The table lists gene names, log2 fold changes for G3BP1 IP versus control IP, associated *P*-values, and −log10(*P*) values. Positive logFC values indicate enrichment in the G3BP1 IP fraction, whereas negative values indicate reduced recovery relative to control IP.

## REFERENCES

1. V. Okur et al., De novo mutations in CSNK2A1 are associated with neurodevelopmental abnormalities and dysmorphic features. Hum Genet 135, 699–705 (2016).

2. N. Belnap et al., Inherited CSNK2A1 variants in families with Okur-Chung neurodevelopmental syndrome. Clin Genet 104, 607–609 (2023).

3. H. Goel, S. O’Donnell, Inherited loss of function variant in CSNK2A1: the oldest reported cases of Okur–Chung syndrome in a single family. Clinical Dysmorphology 33, 121–124 (2024).

4. S. Ramadesikan et al., Expanding the phenotypic spectrum of CSNK2A1-associated Okur-Chung neurodevelopmental syndrome. Human Genetics and Genomics Advances 6, (2025).

5. E. D. Bagatelas, M. M. Khan, G. V. Rushing, OCNDS core features are conserved across variants, with loop-region mutations driving greater symptom burden. Frontiers in Human Neuroscience Volume 19 - 2025, (2025).

6. H. Jafari Khamirani et al., Clinical Features of Okur-Chung Neurodevelopmental Syndrome: Case Report and Literature Review. Mol Syndromol 13, 381–388 (2022).

7. S. Xu, Q. Lian, J. Wu, L. Li, J. Song, Dual molecular diagnosis of tricho-rhino-phalangeal syndrome type I and Okur-Chung neurodevelopmental syndrome in one Chinese patient: a case report. BMC Med Genet 21, 158 (2020).

8. K. Ahmed, D. A. Gerber, C. Cochet, Joining the cell survival squad: an emerging role for protein kinase CK2. Trends in cell biology 12, 226–230 (2002).

9. K. Niefind, B. Guerra, I. Ermakowa, O. G. Issinger, Crystal structure of human protein kinase CK2: insights into basic properties of the CK2 holoenzyme. The EMBO journal 20, 5320–5331 (2001).

10. S. Sarno et al., Basic Residues in the 74–83 and 191–198 Segments of Protein Kinase CK2 Catalytic Subunit are Implicated in Negative but not in Positive Regulation by the β-Subunit. European journal of biochemistry 248, 290–295 (1997).

11. T. Buchou et al., Disruption of the regulatory β subunit of protein kinase CK2 in mice leads to a cell-autonomous defect and early embryonic lethality. Molecular and cellular biology 23, 908–915 (2003).

12. D. Y. Lou et al., The alpha catalytic subunit of protein kinase CK2 is required for mouse embryonic development. Molecular and cellular biology 28, 131–139 (2008).

13. P. Blanquet, Casein kinase 2 as a potentially important enzyme in the nervous system. Progress in neurobiology 60, 211–246 (2000).

14. J. Castello, A. Ragnauth, E. Friedman, H. Rebholz, CK2—an emerging target for neurological and psychiatric disorders. Pharmaceuticals 10, 7 (2017).

15. J. Castello et al., CK2 regulates 5-HT4 receptor signaling and modulates depressive-like behavior. Molecular psychiatry 23, 872–882 (2018).

16. H. Rebholz et al., CK2 negatively regulates Gαs signaling. Proceedings of the National Academy of Sciences 106, 14096–14101 (2009).

17. C. Borgo, C. D’Amore, S. Sarno, M. Salvi, M. Ruzzene, Protein kinase CK2: a potential therapeutic target for diverse human diseases. Signal transduction and targeted therapy 6, 183 (2021).

18. A. F. Rosenberger et al., Increased occurrence of protein kinase CK2 in astrocytes in Alzheimer’s disease pathology. Journal of neuroinflammation 13, 4 (2016).

19. M. Y. Ryu et al., Localization of CKII β subunits in Lewy bodies of Parkinson’s disease. Journal of the neurological sciences 266, 9–12 (2008).

20. L. C. Reineke et al., Casein Kinase 2 Is Linked to Stress Granule Dynamics through Phosphorylation of the Stress Granule Nucleating Protein G3BP1. Mol Cell Biol 37, (2017).

21. K. Sato, K.-i. Takayama, S. Inoue, Stress granules sequester Alzheimer’s disease-associated gene transcripts and regulate disease-related neuronal proteostasis. Aging 15, 3984–4011 (2023).

22. D. M. Caefer et al., The Okur-Chung Neurodevelopmental Syndrome Mutation CK2(K198R) Leads to a Rewiring of Kinase Specificity. Front Mol Biosci 9, 850661 (2022).

23. J. M. Cruz-Gamero et al., Missense mutation in the activation segment of the kinase CK2 models Okur-Chung neurodevelopmental disorder and alters the hippocampal glutamatergic synapse. Mol Psychiatry 30, 1497–1509 (2025).

24. I. Dominguez et al., Okur-Chung neurodevelopmental syndrome-linked CK2alpha variants have reduced kinase activity. Hum Genet, (2021).

25. N. C. Anderson et al., Balancing serendipity and reproducibility: Pluripotent stem cells as experimental systems for intellectual and developmental disorders. Stem Cell Reports 16, 1446–1457 (2021).

26. A. Gast, C. Werner, K. Niefind, J. Jose, Functional characterization of 42 CK2α de novo variants associated with Okur-Chung neurodevelopmental syndrome. The FEBS Journal n/a, (2026).

27. P. K. Sahoo et al., Disruption of G3BP1 granules promotes mammalian CNS and PNS axon regeneration. Proc Natl Acad Sci U S A 122, e2411811122 (2025).

28. P. K. Sahoo et al., A Ca(2+)-Dependent Switch Activates Axonal Casein Kinase 2alpha Translation and Drives G3BP1 Granule Disassembly for Axon Regeneration. Curr Biol 30, 4882–4895 e4886 (2020).

29. P. K. Sahoo et al., Axonal G3BP1 stress granule protein limits axonal mRNA translation and nerve regeneration. Nat Commun 9, 3358 (2018).

30. A. Aulas et al., G3BP1 promotes stress-induced RNA granule interactions to preserve polyadenylated mRNA. J Cell Biol 209, 73–84 (2015).

31. D. M. Parker, D. Tauber, R. Parker, G3BP1 promotes intermolecular RNA-RNA interactions during RNA condensation. Molecular Cell 85, 571–584.e577 (2025).

32. J. L. Twiss, C. N. Buchanan, Regulation of Subcellular Protein Synthesis for Restoring Neural Connectivity. Int J Mol Sci 26, (2025).

33. H. Sidibé, A. Dubinski, C. Vande Velde, The multi-functional RNA-binding protein G3BP1 and its potential implication in neurodegenerative disease. J Neurochem 157, 944–962 (2021).

34. P. K. Sahoo, D. S. Smith, N. Perrone-Bizzozero, J. L. Twiss, Axonal mRNA transport and translation at a glance. J Cell Sci 131, (2018).

35. A. Biever, P. G. Donlin-Asp, E. M. Schuman, Local translation in neuronal processes. Current Opinion in Neurobiology 57, 141–148 (2019).

36. R. Taylor, N. Nikolaou, RNA in axons, dendrites, synapses and beyond. Front Mol Neurosci 17, 1397378 (2024).

37. I. Dalla Costa et al., The functional organization of axonal mRNA transport and translation. Nature Reviews Neuroscience 22, 77–91 (2021).

38. C. E. Holt, K. C. Martin, E. M. Schuman, Local translation in neurons: visualization and function. Nature structural & molecular biology 26, 557–566 (2019).

39. M. Agrawal et al., Restoring DSCAM expression mitigates neuronal morphology and axon guidance deficits in Down syndrome. Brain, (2026).

40. S. Staebler et al., In long-lasting cellular stress phases of melanoma cells, stress granules are dissolved by HSP70. Cellular and Molecular Life Sciences 82, 366 (2025).

41. K. Lavrynenko et al., mRNA recruitment by G3BP1 condensates is regulated by Caprin1 but requires G3BP1 binding to mRNA. Sci Rep 15, 37076 (2025).

42. T. Schulte et al., Caprin-1 binding to the critical stress granule protein G3BP1 is influenced by pH. Open Biol 13, 220369 (2023).

43. J. Guillén-Boixet et al., RNA-Induced Conformational Switching and Clustering of G3BP Drive Stress Granule Assembly by Condensation. Cell 181, 346–361.e317 (2020).

44. H. Matsuki et al., Both G3BP1 and G3BP2 contribute to stress granule formation. Genes Cells 18, 135–146 (2013).

45. M. Desai, K. Gulati, M. Agrawal, S. Ghumra, P. K. Sahoo, Stress granules: Guardians of cellular health and triggers of disease. Neural Regen Res 21, 588–597 (2026).

46. P. Yang et al., G3BP1 Is a Tunable Switch that Triggers Phase Separation to Assemble Stress Granules. Cell 181, 325–345 e328 (2020).

47. F. Meggio, L. A. Pinna, One-thousand-and-one substrates of protein kinase CK2? Faseb j 17, 349–368 (2003).

48. J. O. Bush, P. Soriano, Ephrin-B1 regulates axon guidance by reverse signaling through a PDZ-dependent mechanism. Genes Dev 23, 1586–1599 (2009).

49. Y. Z. Alabed, E. Grados-Munro, G. B. Ferraro, S. H. K. Hsieh, A. E. Fournier, Neuronal responses to myelin are mediated by rho kinase. Journal of Neurochemistry 96, 1616–1625 (2006).

50. N. Arimura et al., Phosphorylation by Rho Kinase Regulates CRMP-2 Activity in Growth Cones. Molecular and Cellular Biology 25, 9973–9984 (2005).

51. J. F. Borisoff et al., Suppression of Rho-kinase activity promotes axonal growth on inhibitory CNS substrates. Molecular and Cellular Neuroscience 22, 405–416 (2003).

52. Y. Fujita, T. Yamashita, Axon growth inhibition by RhoA/ROCK in the central nervous system. Frontiers in Neuroscience Volume 8 - 2014, (2014).

53. C. H. He et al., Overexpression of EphB6 and EphrinB2 controls soma spacing of cortical neurons in a mutual inhibitory way. Cell Death Dis 14, 309 (2023).

54. D. S. Kang et al., Netrin-1/DCC-mediated PLCγ1 activation is required for axon guidance and brain structure development. The EMBO Reports 19, EMBR201846250 (2018).

55. M. Kinoshita-Kawada et al., A crucial role for Arf6 in the response of commissural axons to Slit. Development 146, (2019).

56. J. Mills et al., Role of integrin-linked kinase in nerve growth factor-stimulated neurite outgrowth. J Neurosci 23, 1638–1648 (2003).

57. S. S. Parker et al., Competing molecular interactions of aPKC isoforms regulate neuronal polarity. Proc Natl Acad Sci U S A 110, 14450–14455 (2013).

58. B. Rico et al., Control of axonal branching and synapse formation by focal adhesion kinase. Nat Neurosci 7, 1059–1069 (2004).

59. B. Torroba, A. Herrera, A. Menendez, S. Pons, PI3K regulates intraepithelial cell positioning through Rho GTP-ases in the developing neural tube. Dev Biol 436, 42–54 (2018).

60. J. Zhong et al., Raf kinase signaling functions in sensory neuron differentiation and axon growth in vivo. Nature Neuroscience 10, 598–607 (2007).

61. K. Y. Wu et al., Local translation of RhoA regulates growth cone collapse. Nature 436, 1020–1024 (2005).

62. S. Mollet et al., Translationally repressed mRNA transiently cycles through stress granules during stress. Mol Biol Cell 19, 4469–4479 (2008).

63. J. R. Buchan, R. Parker, Eukaryotic stress granules: the ins and outs of translation. Mol Cell 36, 932–941 (2009).

64. M. W. Antoine, Paradoxical Hyperexcitability in Disorders of Neurodevelopment. Front Mol Neurosci 15, 826679 (2022).

65. T. A. Fenton et al., Hyperexcitability and translational phenotypes in a preclinical mouse model of SYNGAP1-related intellectual disability. Translational Psychiatry 14, 405 (2024).

66. Y. Hussein et al., Early maturation and hyperexcitability is a shared phenotype of cortical neurons derived from different ASD-associated mutations. Translational Psychiatry 13, 246 (2023).

67. Y. Takarae, J. Sweeney, Neural Hyperexcitability in Autism Spectrum Disorders. Brain Sci 7, (2017).

68. D. Ballardin, J. M. Cruz-Gamero, T. Bienvenu, H. Rebholz, Comparing Two Neurodevelopmental Disorders Linked to CK2: Okur-Chung Neurodevelopmental Syndrome and Poirier-Bienvenu Neurodevelopmental Syndrome-Two Sides of the Same Coin? Front Mol Biosci 9, 850559 (2022).

69. M. Nakashima et al., Identification of de novo CSNK2A1 and CSNK2B variants in cases of global developmental delay with seizures. Journal of Human Genetics 64, 313–322 (2019).

70. M. Agrawal, K. Welshhans, Local Translation Across Neural Development: A Focus on Radial Glial Cells, Axons, and Synaptogenesis. Frontiers in Molecular Neuroscience Volume 14 - 2021, (2021).

71. A.-S. Hafner, P. G. Donlin-Asp, B. Leitch, E. Herzog, E. M. Schuman, Local protein synthesis is a ubiquitous feature of neuronal pre- and postsynaptic compartments. Science 364, eaau3644 (2019).

72. D. Rajgor, T. M. Welle, K. R. Smith, The Coordination of Local Translation, Membranous Organelle Trafficking, and Synaptic Plasticity in Neurons. Front Cell Dev Biol 9, 711446 (2021).

73. E. H. Pool et al., Aberrant phase separation of two PKA RIβ neurological disorder mutants leads to mechanistically distinct signaling deficits. Cell Rep 44, 115797 (2025).

74. N. B. Illarionova, M. P. Moshkin, The Role of RNA-protein Granules in Neurodevelopment. Biology Bulletin Reviews 16, 384–393 (2026).

75. N. Bencsik, D. Kimsanaliev, K. Tárnok, K. Schlett, Stress-Induced Membraneless Organelles in Neurons: Bridging Liquid–Liquid Phase Separation and Neurodevelopmental Dysfunction. International Journal of Molecular Sciences 26, 9068 (2025).

76. B. T. Hurtle, L. Xie, C. J. Donnelly, Disrupting pathologic phase transitions in neurodegeneration. The Journal of Clinical Investigation 133, (2023).

77. A. Naskar, A. Nayak, M. R. Salaikumaran, S. S. Vishal, P. P. Gopal, Phase separation and pathologic transitions of RNP condensates in neurons: implications for amyotrophic lateral sclerosis, frontotemporal dementia and other neurodegenerative disorders. Frontiers in Molecular Neuroscience Volume 16 - 2023, (2023).

78. R. Nóbrega-Martins et al., RNA Granules at the Crossroads of Synaptic Dysfunction and Neurodegeneration. J Neurochem 169, e70269 (2025).

79. S. P. Mojica-Perez et al., Protocol for selecting single human pluripotent stem cells using a modified micropipetter. STAR protocols 4, (2023).

80. A. M. Tidball et al., Genome-wide CRISPRi Screen in Human iNeurons to Identify Novel Focal Cortical Dysplasia Genes. bioRxiv, (2023).

81. A. M. Tidball et al., Deriving early single-rosette brain organoids from human pluripotent stem cells. Stem Cell Reports 18, 2498–2514 (2023).

82. L. Kershner, K. Welshhans, RACK1 is necessary for the formation of point contacts and regulates axon growth. Dev Neurobiol 77, 1038–1056 (2017).

83. S. Ghumra, M. Agrawal, M. Desai, L. Kuchibatla, P. K. Sahoo, Isolation and Quantification of Axonal mRNAs Using Porous Membrane Inserts and RTddPCR. J Vis Exp, (2026).

84. T. T. Merianda, D. Vuppalanchi, S. Yoo, A. Blesch, J. L. Twiss, Axonal transport of neural membrane protein 35 mRNA increases axon growth. Journal of Cell Science 126, 90–102 (2013).

85. K. M. O’Neill et al., Time-dependent homeostatic mechanisms underlie brain-derived neurotrophic factor action on neural circuitry. Communications Biology 6, 1278 (2023).

86. A. R. Rodríguez, K. M. O’Neill, P. Swiatkowski, M. V. Patel, B. L. Firestein, Overexpression of cypin alters dendrite morphology, single neuron activity, and network properties via distinct mechanisms. Journal of neural engineering 15, 016020 (2018).

87. R. Q. Quiroga, Z. Nadasdy, Y. Ben-Shaul, Unsupervised spike detection and sorting with wavelets and superparamagnetic clustering. Neural computation 16, 1661–1687 (2004).

88. I. Rishal et al., WIS-NeuroMath enables versatile high throughput analyses of neuronal processes. Dev Neurobiol 73, 247–256 (2013).

